# SWI/SNF chromatin remodeling determines brassinosteroid-induced transcriptional activation

**DOI:** 10.1101/2023.06.23.544932

**Authors:** Tao Zhu, Chuangqi Wei, Yaoguang Yu, Jiameng Zhu, Zhenwei Liang, Yuhai Cui, Zhi-Yong Wang, Chenlong Li

## Abstract

The brassinosteroid (BR) hormone is a central modulator of plant growth, development, and responses to stresses by activating or repressing the expression of thousands of genes through the transcription factor BRASSINAZOLE-RESISTANT 1 (BZR1) and its homologues. However, the molecular mechanism that determines the transcriptional activation versus repression activity of BZR1 remains largely unclear. Here, we show that BZR1-responsive transcriptional activation at thousands of loci requires the Switch defective/sucrose non-fermentable (SWI/SNF)-complexes-mediated chromatin accessibility regulation. BR-activated BZR1 controls the activation or repression of thousands of BZR1 target genes through reprograming genome-wide chromatin accessibility landscape in *Arabidopsis thaliana*. BZR1 physically interacts with the <u>B</u>RAHMA (BRM)-<u>A</u>ssociated <u>S</u>WI/SNF complexes (BAS), co-localizes with BRM on the genome, and enhances BRM occupancy at sites of increased accessibility by BR. Loss of BRM abrogates the capacity of BZR1 to increase but not decrease chromatin accessibility, blocks BR-induced hypocotyl elongation, and diminishes BZR1-mediated transcriptional activation rather than repression. Together, our work reveals that the BAS chromatin remodeling complex is a critical epigenetic regulatory partner in dictating BZR1-mediated transcriptional activation ability, thus providing a long sought mechanistic explanation for how BR signaling activates gene transcription in shaping diverse developmental programs.

**Teaser:** BZR1-responsive transcriptional activation activity at thousands of loci requires the SWI/SNF-complexes-mediated chromatin accessibility regulation.

## Introduction

To ensure survival, plants must sense and respond to various environmental signals (*1*). Thus, plants need efficient ways to communicate between cells and cooperate within tissues in the developmental responses to external signals. Many signaling molecules are used to accomplish this process. One such class of signaling molecules is BR, a polyhydroxylated steroidal hormone involved in diverse growth and development processes (*2, 3*). BR is recognized by the extracellular leucine-rich repeat (LRR) domains of cell transmembrane receptor kinase BRASSINOSTEROID-INSENSITIVE 1 (BRI1). BR binding enhances BRI1 heteromerization with BRI1-associated kinase 1 (BAK1) (*4, 5*), which triggers a series of phosphorylation events and the activation of BR-SIGNALING KINASE 1 (BSK1) and CONSTITUTIVE DIFFERENTIAL GROWTH 1 (CDG1) (*6, 7*). BSK1 and CDG1 further activates BRI-SUPPRESSOR 1 (BSU1) family phosphatases (*7*). The activated BSU1 dephosphorylates and inactivates the primary negative regulator GLYCOGEN SYNTHASE KINASE 3 (GSK3)-like kinase BRASSINOSTEROID-INSENSITIVE 2 (BIN2) (*8*), leading to the dephosphorylation and activation of master transcription factors BRASSINAOLE-RESISTANT 1 (BZR1) and BRI-EMS-SUPPRESSOR 1 (BES1) by PROTEIN PHOSPHATASE 2A (PP2A) (*9*). The dephosphorylated BZR1 and BES1 are transported to the nucleus to achieve transcriptional regulation of thousands of BR responsive genes.

BZR1 is atypical basic helix–loop–helix (bHLH) transcription factor that functions in orchestrating diverse developmental and physiological processes (*2, 3, 10*). For example, BZR1 is an essential component of the transcriptional activation module that regulates hypocotyl elongation in response to light, temperature, auxin, gibberellin, and sugar (*11–14*). The gradient of BZR1 activity in the root tip controls the balance of stem cell maintenance and differentiation in the root meristem (*15*), whereas its activities in specific cell types regulates xylem differentiation, cell division, and symbiosis (*16, 17*). BZR1-mediated transcriptional activation is also required for fertility and plays specific roles in the development of anther, pollen, and seed (*18, 19*). In the shoot meristem, BZR1 represses organ boundary identity genes to regulate shoot architecture (*20*). Beyond their roles in growth and development, BZR1 plays roles in regulating immune responses and balancing the trade-off between growth and immunity (*21–24*). There is also evidence that BZR1 is involved in acclimation to heat, cold, and drought stresses (*10, 11, 25, 26*). These observations underscore the essentiality of transcriptional regulation mediated by BR-BZR1 signaling in the context of plant growth, development, and immune responses.

Although it has been known for decades that BZR1 and BES1 transcription factors can either activate or repress BR-responsive genes (*10*), the mechanistic basis of this dichotomy is still poorly understood. Current evidence suggests that different cis elements might be related to the activation and repression ability of BZR1. Indeed, earlier studies showed that BZR1-induced genes enrich E-box (CANNTG) motif, whereas BZR1-repressed genes enrich BR-response elements (BRRE, CGTG(T/C)G) (*11, 27, 28*). Intriguingly, two nucleobases flanking the core binding G-box (CACGTG) motif were recently proposed by analysis of in vitro DNA affinity purification sequencing (DAP-seq) data to be responsible for BZR1-responsive transcriptional repression rather than transcriptional activation (*29*). However, whether and how *cis* elements may contribute to distinguish transcriptional activation from repression activity of BZR1 remain undetermined. Interestingly, BZR1 was shown to interact with transcriptional co-repressor TOPLESS (TPL) family proteins through its ERF associated amphiphilic repression (EAR) domain and recruits TPL to BZR1-repressed genes (*30*). The BZR1-TPL complex further allows the recruitment of Histone deacetylase 19 (HDA19) to mediate histone deacetylation and thus mediates BR responsive gene repression (*30*). Notably, BZR1 has been reported to activate several cell elongation-related genes by linking to the PICKLE chromatin remodeler (*31*). Additionally, BZR1 was shown to induce floral repressor *FLOWERING LOCUS C* (*FLC*) in connection with the histone 3 lysine 27 (H3K27) demethylase EARLY FLOWERING 6 (*32*). Transcription factors PHYTOCHROME INTERACTING FACTOR 4 (PIF4), BLUE-LIGHT INHIBITOR OF CRYPTOCHROMES1 (BIC1), and AUXIN RESPSONE FACTOR 6 (ARF6) were shown to interact with BZR1 and cooperatively up-regulate genes involved in cell elongation in response to light and temperature signaling (*11, 13, 33*). However, the fundamental principle determining BZR1-mediated transcriptional activation is still largely unknown.

Chromatin accessibility is crucial in regulating gene expression and has a dynamic response to endogenous and exogenous signals (*34*). In eukaryotes, the adenosine triphosphate (ATP)-dependent chromatin-remodeling enzymes disrupt histone contacts and translocate DNA around the nucleosome to slide, evict, exchange or assemble the histone octamer, thus regulating the accessibility to DNA (*35–37*). Switch defective/sucrose non-fermentable (SWI/SNF) complexes are chromatin remodelers responsible for increasing DNA accessibility (*38*). Active regulatory DNA regions require continuous chromatin remodeling activity of SWI/SNF complexes; therefore, their activities must be tightly regulated to ensure fidelity and plasticity of genomic processes (*39, 40*). In *Arabidopsis*, three subclasses of SWI/SNF chromatin remodeling complexes were identified: BRM-associated SWI/SNF complexes (BAS), SPLAYED associated SWI/SNF complexes (SAS), and MINUSCULE-associated SWI/SNF complexes (MAS) (*41, 42*). However, the molecular mechanisms responsible for the precise localization of SWI/SNF chromatin remodeling complexes to specific genomic loci, therefore ensuring their proper activity during the intricate processes of growth and development, remain obscure. Although both SWI/SNF and BR are vital for diverse plant developmental processes, no direct molecular connection has been established between SWI/SNF-mediated genome accessibility regulation and BR-directed dynamic hormone signaling network during development.

In this study, we demonstrate that BAS-type SWI/SNF complexes are required for BR signaling to mediate chromatin accessibility landscape of thousands of loci to dictate BR-responsive transcriptional activation. We show that BZR1 physically interacts with BAS-complexes subunits and co-localizes extensively with BAS on the genome, with higher BRM enrichment at sites where BR increases chromatin accessibility. BR signaling enhances BRM occupancy at BZR1-increased accessible loci. Loss of BRM nearly completely abolishes BZR1-mediated increase rather than decrease, of chromatin accessibility. Consistently, genetic disruption of BRM blocks BZR1-mediated hypocotyl elongation in the dark and gene transcriptional activation but not repression activity, highlighting SWI/SNF chromatin remodeler complexes as specific and critical regulators of BR-mediated gene activation. In summary, our findings unravel that the BAS chromatin remodeling complex is a critical epigenetic regulatory partner that determines the transcriptional activation activity of BZR1 in the BR signaling pathway. Our work also sheds light on hormone information in directing global epigenome activation, with broad relevance for the developmental control of plants.

## Results

### BR-BZR1 signaling modulates chromatin accessibility landscape

To explore how BR signaling might regulate the chromatin accessibility landscape, we harvested Col wild-type (WT) and *bri1-701* mutant Arabidopsis seedlings grown in the dark for five days and performed assay for transposase-accessible chromatin by sequencing (ATAC-seq). We identified 2,658 differentially accessible regions (DARs, |log_2_ fold change| ≥ 0.4) between Col and *bri1-701*, of which 57% and 43% showed decreased and increased accessibility, respectively, in *bri1-701* mutants (Fig. 1, A to D and fig. S1, A and B and table S1). DARs were predominantly located in regions near the transcription initiation sites (TSSs) of genes (Fig. 1E). These results suggest that BR has a dual function in regulating TSS chromatin accessibility, probably underscoring the dual role of BR to activate and repress gene transcription.

**Fig. 1.**
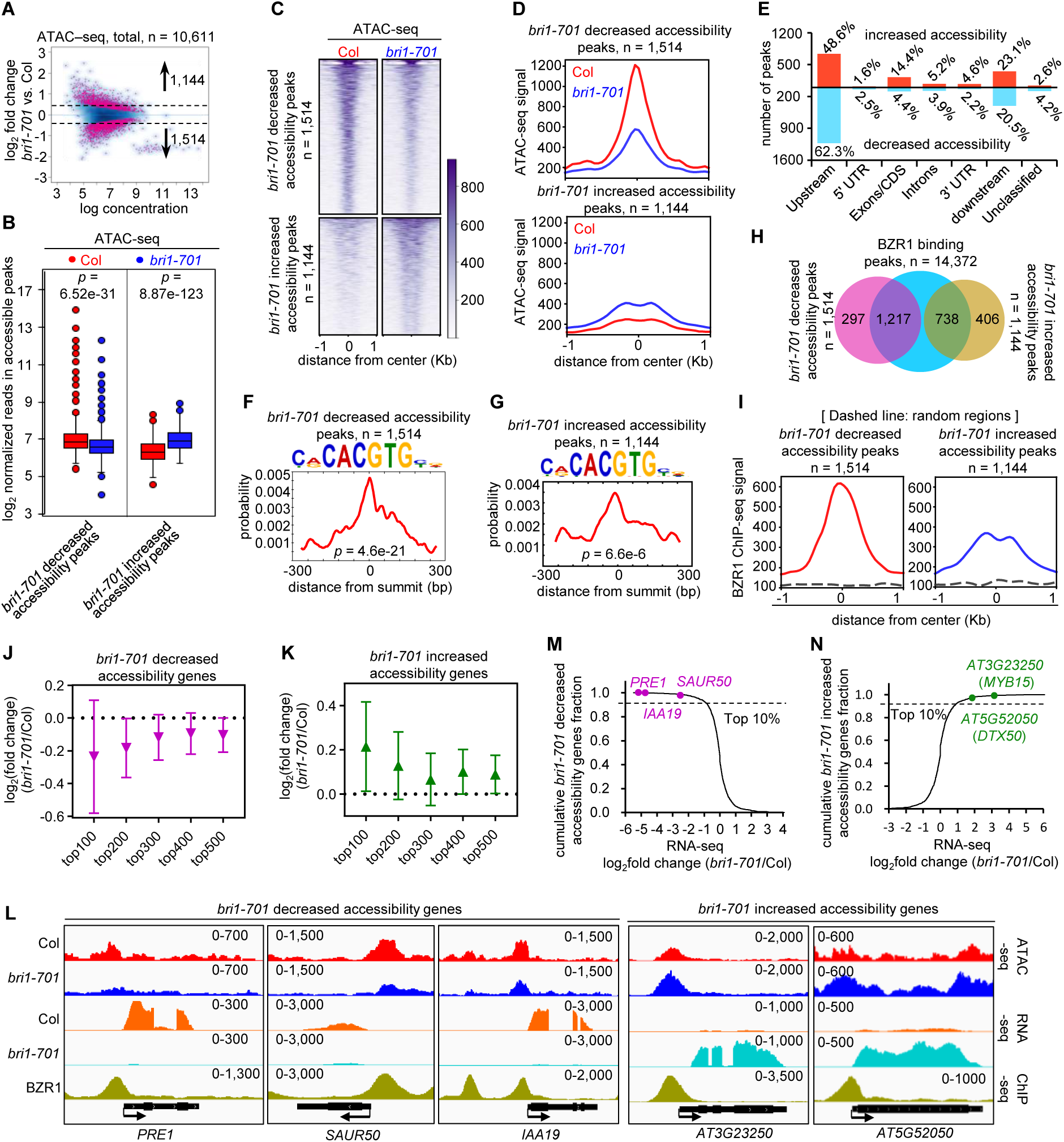
BR limitation induces the genome-wide changes of chromatin accessibility landscape. (**A**) Scatter plot showing fold-change (|log_2_ fold change| ≥ 0.4) of accessible peaks between WT and *bri1-701*. Blue dots, stable peaks; pink dots, differential peaks. The numbers of differentially accessible peaks (increased or decreased) according to FDR < 0.05. (**B**) Box plots showing counts at regions that decreased accessibility and increased accessibility in *bri1-701* for the indicated ATAC-seq experiments. (**C**), (**D**) Heatmap (C) and metagene plots (D) reflecting the ATAC-seq signals over the *bri1-701* decreased, or increased chromatin accessibility sites for the indicated ATAC-seq experiments. (**E**) Bar chart showing the distribution of changed chromatin accessible peaks at genic and intergenic regions in the genome. (**F**), (**G**) The G-Box element is significantly enriched in *bri1-701* decreased or increased accessibility peaks. (**H**) Venn diagram showing statistically significant overlaps between the BR-regulated accessibility peaks and the BZR1 binding peaks. (**I**) Metagene plots reflecting the occupancy of BZR1 over the *bri1-701* decreased, or increased chromatin accessibility sites (**J**), (**K**) The correlation between the magnitude of the changes in the *bri1-701* mutants for chromatin accessibility and gene expression. (**L**) IGV view of ATAC-seq, RNA-seq and ChIP-seq of indicated samples at the *bri1-701* decreased or increased accessibility genes. The black diagrams underneath indicate gene structure. The y-axis scales represent shifted merged MACS2 tag counts for every 10-bp window. (**M**) Cumulative distribution function plot reflecting *bri1-701* down-regulated expression genes in *bri1-70*1 decreased accessibility genes, the top one-tenth fraction reflects genes associated with the top changed genes. (**N**) Cumulative distribution function plot reflecting *bri1-701* up-regulated expression genes in *bri1-70*1 increased accessibility genes, the top one-tenth fraction reflects genes associated with the top changed genes.

Next, we wondered whether these changes in chromatin accessibility were directly regulated by BZR1. Using the CentriMo motif analysis pipeline (*43*), we found that BR-regulated chromatin accessibility regions significantly enriched for sequences containing the core G-box (CACGTG) motif (fig. S1C). Further analysis of regions with decreased or increased accessibility in the *bri1-701* mutants showed that G-boxes recognized by BZR1 significantly enriched within both groups (Fig. 1, F and G). We then carried out chromatin immunoprecipitation followed by high-throughput sequencing (ChIP-seq) using Arabidopsis transgenic lines expressing yellow fluorescence protein (YFP)-tagged BZR1 under the control of its native promoter in the Col background (*ProBZR1:BZR1-YFP*) to identify BZR1-enriched genes in the 5-day-old seedlings grown in the dark conditions (table S2). Consistent with the previous ChIP-seq data for BZR1, the known BZR1 target genes were observed in our dataset (fig. S1, D and E). We found that 80.3% of the decreased DARs in *bri1-701* (1,217 out of 1,514 peaks) overlapped with the BZR1-binding regions (Fig. 1H). Significant overlap between the increased DARs in *bri1-701* and the BZR1-binding regions (738 out of 1144 peaks, 64.5%) was also observed (Fig. 1H). Consistently, a highly significant enrichment in ChIP-seq signals for BZR1 was showed at the centers of the increased or decreased DARs (Fig. 1I). Furthermore, 60% of BZR1-targeted DAR them had reduced accessibility, while 40% showed increased accessibility in the *bri1-701* mutants (fig. S1F). Together, these results support the direct role of BZR1 both in increasing and decreasing chromatin accessibility in plants.

We next assessed whether changes in chromatin accessibility caused by BR signaling-deficiency are correlated with changes in expression. Transcriptome profiling by RNA-seq identified a total of 1,729 genes (|log2 fold change| ≥ 1) that were dysregulated in the *bri1-701* mutants, of which 875 and 854 showed down-regulated and up-regulated, respectively (fig. S1G and table S3). We found that decreased and increased DAR genes were significant enriched among genes with decreased and increased expression in the *bri1-701* mutants, respectively (fig. S1H). Moreover, the transcription of the decreased and increased DAR genes was down-regulated and up regulated, respectively, in the *bri1-701* mutants (fig. S1I). We further divided the top 500 genes that showed dysregulated chromatin accessibility in the *bri1-701* mutants into five fractions according to the degree of the change and analyzed the corresponding changes in RNA expression. We found a positive correlation between the magnitude of changes in the *bri1-701* mutants for chromatin accessibility and gene expression (Fig. 1, J and K). These positive correlations were exemplified at individual loci (Fig. 1L). Thus, BR signaling pathway can activate and repress gene expression in a chromatin accessibility-dependent manner.

To understand the potential physiological significance of BR-mediated changes in chromatin accessibility, we identified among the reduced DAR genes the most highly down-regulated genes in the *bri1-701* mutants. The top-regulated genes were those previously found to mediate cell-elongation, including *PACLOBUTRAZOL RESISTANCE 1* (*PRE1*), *SMALL AUXIN UPREGULATED RNA 50* (*SAUR50*), and *INDOLE-3-ACETIC ACID INDUCIBLE 19* (*IAA19*) (*11*) (Fig. 1M), thus suggesting that enhanced chromatin accessibility by BR signaling facilitates transcriptional activation of the cell elongation processes. Gene Ontology (GO) analysis using genes showing decreased accessibility and expression in the *bri1-701* mutants revealed terms related to response to Auxin, light intensity, red or far-red light, and cell-wall organization processes (fig. S1J). In contrast, when we identified the most highly up regulated genes in the *bri1-701* mutants among the increased DAR genes, stress-related genes such as *MYB DOMAIN PROTEIN 15* (*MYB15*) (*44*) and *DETOXIFICATION EFFLUX CARRIER 50* (*DTX50*) (*45*) were observed (Fig. 1N). GO analysis of genes with increased chromatin accessibility and transcription in the *bri1-701* mutants showed a marked excess of terms related to cellular response to hypoxia, salicylic acid, salt stress, and oxidative stress (fig. S1K). This analysis suggests that BR-mediated chromatin accessibility decrease and associated transcriptional down-regulation are involved in stress-responsive processes. Taken together, these results imply that BR maintained genome-wide chromatin accessibility landscape regulates a gene expression axis that may balance plant growth and stress response processes.

### BZR1 interacts with the BAS complexes in plants

We next sought to define the molecular mechanisms by which BR signaling regulates chromatin accessibility. We conducted immunoprecipitation followed by Mass spectrometry (IP/MS) using our previously described *BZR1-YFP* line and identified proteins that co-purified with BZR1 by mass spectrometry. Along with the known BZR1-interacting protein TPL (*30*), we identified the SWI/SNF chromatin remodeler ATPase BRM that co-purified with BZR1 (Fig. 2A). Hemagglutinin (HA)-tagged BZR1 co-immunoprecipitated with FLAG-tagged BRM in *Nicotiana benthamiana* leaves (Fig. 2B). Consistent with the overexpression data, the interaction between BZR1 and BRM was also detected in an Arabidopsis line expressing the BZR1-3FLAG and BRM-GFP proteins under their respective native promoters (Fig. 2C). Recent studies showed that Arabidopsis SWI/SNF complexes can be divided into three types of subcomplexes, including the BRM-Associated SWI/SNF complexes (BAS) (*41, 42*) . BAS complexes contain a series of BAS-subcomplex-specific subunits, including BRAHMA-INTERACTING PROTEINS 1/2 (BRIP1/2), BROMODOMAIN-CONTAINING PROTEIN 2/13 (BRD2/13), and SWI/SNF ASSOCIATED PROTEIN 73A (SWP73A). To further evaluate whether BZR1 forms a complex with BAS, co-immunoprecipitation (Co-IP) assays were performed to detect the interaction between BZR1 and the BAS complex-specific subunits. HA-tagged BRIP1/2, BRD2/13, or SWP73A co immunoprecipitated with FLAG-tagged BZR1 in *N. benthamiana* leaves (Fig. 2, D to H). In addition, bimolecular fluorescence complementation (BiFC) assays using *N. benthamiana* leaves detected positive fluorescent signals in nuclei when co-expressing N-terminal YFP-fused BZR1 and C-terminal YFP-fused BRM or the known BAS specific subunits (fig. S2, A and B). Together, these results demonstrate the tethering of BZR1 to the BAS complexes to form a BZR1-containing BAS complex.

**Fig. 2.**
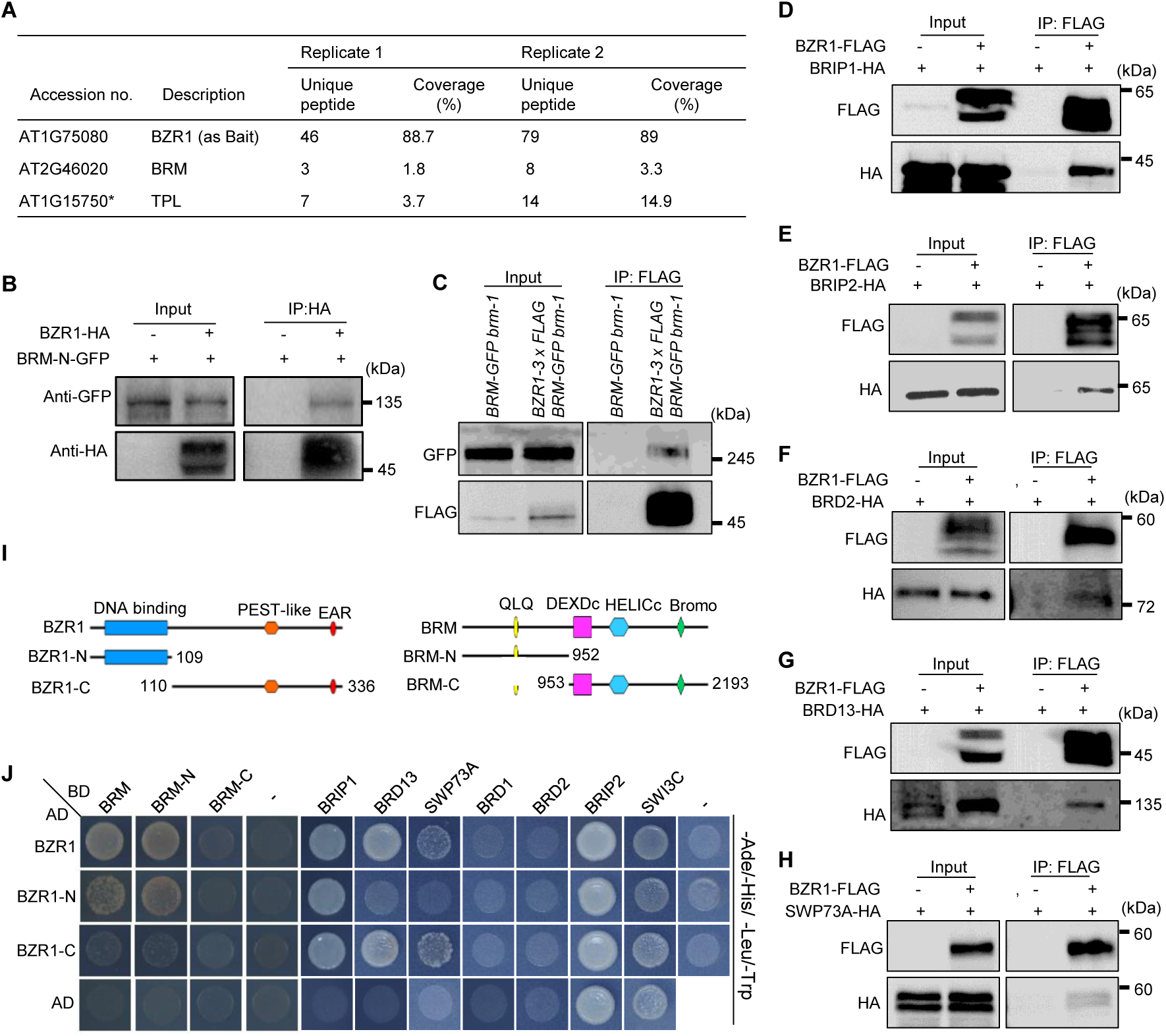
The BZR1 interacts with BAS complex. (**A**) Summary of the peptides of BRM identified by mass spectrometry from an anti GFP purification of a *35S:BZR1-YFP* overexpressed line. Two biological replicates are shown. * represents the known BZR1-interacting protein TPL. (**B**) Co-IP showing the interaction of BZR1 with BRM-N terminal (1-952 amino acids). BRM-N-GFP was coimmunoprecipitated with anti-HA-agarose beads from *Nicotiana benthamiana* leaves that co-expressed BRM-N-GFP and BZR1-HA. (**C**) Immunoblot showing the levels of BRM-GFP and BZR1-3xFLAG from co-IP experiments with anti-FLAG antibody in the genetic backgrounds indicated above lanes. For each plot the antibody used is indicated on the left, and the sizes of the protein markers are indicated on the right. (**D**) to (**H**) Co-IP assays showing the interaction of BZR1 with BRIP1/2, BRD2/13, and SWP73A. BZR1 was coimmunoprecipitated with anti-FLAG-agarose beads from *Nicotiana benthamiana* leaves that co-expressed BRIP1/2-HA, BRD2/13-HA, SWP73A-HA and BZR1-FLAG. (**I**) Schematic illustration of the BZR1 and BRM protein and its truncated versions. (**J**) Yeast two-hybrid assays to examine BZR1 interact with BRM and core members of BAS complex. Yeast cells transformed with the indicated plasmids were plated onto quadruple dropout (Selective) (SC-Ade, - His,-Leu, -Trp) medium. AD, Activation Domain; BD, Binding Domain.

We next carried out yeast two-hybrid (Y2H) assays to determine how BZR1 might directly tether with the BAS complexes. These analyzes indicated that the N-terminal part of BRM (amino acids 1–952) is responsible for the direct interaction with the N-terminal domain of BZR1 (amino acids 1–109) (Fig. 2, I and J and fig. S2C). The N-terminal region of BZR1 has been shown to mediate the protein-protein interaction of BZR1 with numerous proteins (*46*). In addition, Y2H assays also showed that the BZR1 could also directly interact with BAS-specific subunits BRIP1, BRD13 and SWP73A (Fig. 2J). Strikingly, BZR1 did not use the N-terminal domain but instead interacts with these BAS specific subunits through its C-terminal region containing the EAR domain (Fig. 2J). Further deletion analysis revealed that the EAR domain was responsible and sufficient for the interaction between BZR1 and SWP73A subunit (fig. S2, D and E). Taken together, these data suggest that BZR1 assembles into the BAS complexes through at least two mechanisms: the N-terminal domain mediates its interaction with the BRM ATPase, and its C-terminal region containing the EAR domain interacts with core module subunits including SWP73A.

### BRM co-location with BZR1 on chromatin

We subsequently assessed the potential genomic interplay between BZR1 and BRM. We carried out the ChIP-seq assay using our previously reported *ProBRM:BRM-GFP brm-1* plants to identify BRM-occupied genes in 5-day-old seedlings grown in the dark. The sets of genes enriched by BRM (table S2) and those by BZR1 exhibited significant overlap, with 65% of the BRM-occupied genes also enriched by BZR1 (Fig. 3, A and B). Furthermore, the distribution patterns of BRM peaks over gene units and flanking intergenic regions were similar to those of BZR1 (fig. S3A), with the strongest enrichment around the TSSs of target genes (fig. S3, B and C). Of note, the G-box-like motif was the top-ranked DNA motifs enriched in BZR1-BRM co-binding sites (fig. S3D). Correlation analysis with ChIP-seq signals for BZR1 confirmed positively correlated BZR1 (*r* = 0.71) co-localization with BRM (Fig. 3C). When we performed correlation analysis of ChIP-seq signal between BZR1 and BAS-specific subunits BRIP1/2 and BRD1/2/13 using our published ChIP-seq data (*47, 48*), we found that BZR1 also showed significantly correlated co-localization with these BAS-specific subunits (Fig. 3C). Consistently, heatmap analysis at BZR1 or BRM binding peaks showed similar enrichment patterns for BZR1, BRM, BRIP1/2, and BRD1/2/13 when we ranked the peaks by BZR1 or BRM signal, respectively (Fig. 3D, E). When we repeated the co-occupancy analysis using enrichment relative to TSSs rather than binding peaks, we found that BZR1-enriched TSSs were also substantially occupied by BRM and those BAS-specific subunits BRIP1/2 and BRD1/2/13 (fig. S3, E and F). We further compared BZR1-bound peaks (*n* = 14,372) with randomly selected BZR1 unbound regions (*n* = 14,372), finding a significant enrichment of BRM at BZR1-bound versus BZR1-unbound regions (Fig. 3F). Similarly, a strong enrichment of BZR1 was observed at BRM-bound (*n* = 15,565) versus BRM-unbound regions (*n* = 15,565) (Fig. 3G).

**Fig. 3.**
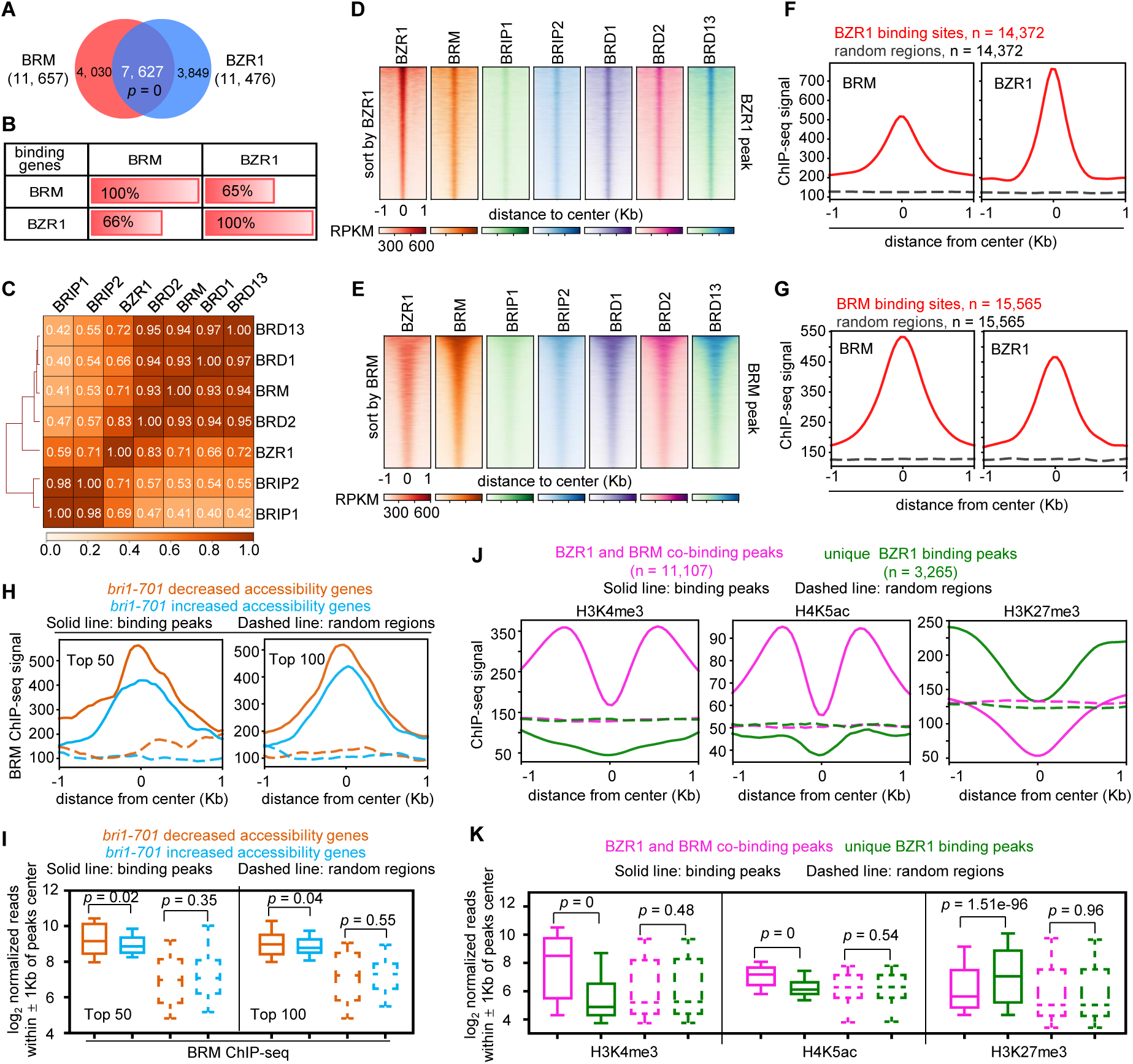
BZR1 co-localizes with BRM genome wide. (**A**) Venn diagrams displaying statistically significant overlaps among genes occupied by BRM and BZR1. The numbers in brackets indicate the total number of genes occupied by BRM, BZR1. *p* values were calculated by the hypergeometric test. (**B**) Percentages of BRM, BZR1 binding genes (by row) overlapping with other binding genes (by column). Shading indicates the strength of overlap. (**C**) Matrix depicting Person correlation coefficients between ChIP–seq datasets, calculated using the bin mode (bin size = 1,000). (**D**) Heatmap representations of ChIP–seq of BZR1, BRM, BRIP1/2, and BRD1/2/3. Rank order is from highest to lowest BZR1-binding peaks signal. log_2_ enrichment was normalized to reads per genome coverage. Read counts per gene were averaged in 50-nucleotide (nt) bins. (**E**) Heatmap representations of ChIP– seq of BZR1, BRM, BRIP1/2, and BRD1/2/3. Rank order is from highest to lowest BRM-binding peaks signal. log_2_ enrichment was normalized to reads per genome coverage. Read counts per gene were averaged in 50-nucleotide (nt) bins. (**F**) Metagene plots displaying the ChIP-seq signals of BRM at BZR1 binding peaks. (**G**) Metagene plots displaying the ChIP-seq signals of BZR1 at BRM binding peaks. (**H**) Metagene plots displaying the ChIP-seq signals of BRM binding peaks at 50 genes (top 50) or 100 genes (top 100) showing decreased or increased accessibility in the *bri1-701* mutants. (**I**) Box plots displaying read counts at *bri1-701* decreased or increased accessibility genes for the BRM ChIP-seq data. Reads were summed ± 1 Kb from the peak center. Significance analysis was determined by two tailed Mann-Whitney U test. (**J**) Metagene plots displaying the ChIP-seq signals of H3K4me3, H4K5ac, and H3K27me3 at BZR1 and BRM co-binding peaks or unique BZR1 binding peaks. (**K**) Box plots displaying read counts at BZR1 and BRM co-binding peaks or unique BZR1 binding peaks for the H3K4me3, H4K5ac, and H3K27me3 ChIP-seq data. Reads were summed ± 1 kb from the peak center. Significance analysis was determined by two tailed Mann-Whitney U test.

Further, we explored the enrichment levels of BRM at *bri1-701* decreased and increased chromatin accessibility sites, observing a higher enrichment of BRM at *bri1-701* decreased chromatin accessibility sites (top 50 and top 100) compared with *bri1-701* increased chromatin accessibility sites (top 50 and top 100) (Fig. 3, H and I). A similar trend held when we repeated the analysis using enrichment relative to TSSs (fig. S3, G and H). These results indicate that BR-dependent chromatin accessibility increased sites have higher BRM occupancy than BR-dependent chromatin accessibility decreased sites.

We also compared the enrichment levels of histone modifications between BZR1-BRM co-binding sites and unique BZR1-binding sites. We found that BZR1-BRM co-binding sites exhibited higher levels of activate histone modification markers, including H3K4me3, H4K5ac, H3K9ac, H3K27ac, H4K8ac, H4K12ac, H4K16ac, H3K4me2, and H3K36me3) compared with BZR1-unique binding sites (Fig. 3, J and K and fig. S4, A and C). On the contrary, BZR1-BRM co-binding sites had lower levels of repressive marker (H3K27me3) compared with BZR1 unique binding sites (Fig. 3, J and K and fig. S4C). Consistently, BZR1-BRM co-binding sites displayed a stronger Pol II enrichment relative to BZR1 unique binding sites (fig. S4, A and B). Together, these results suggest that the physical presence of BRM at the BZR1-BRM co-bound sites may prepare an active chromatin landscape for BR-mediated transcriptional activation.

### BRs enhance BRM targeting at *bri1-701* decreased accessibility sites

Because of the higher BRM enrichment at sites showing decreased accessibility in *bri1-701* mutants compared with sites showing increased chromatin accessibility sites, we sought to assess the potential role of BZR1 in enhancing BAS complexes occupancy to these two chromatin regions. To this end, we used Propiconazole (PPZ), a potent BR biosynthesis inhibitor that inhibits the dephosphorylation of BZR1 and thereby preventing it from entering the nucleus (*49*). As expected, PPZ-treated plants showed shortened hypocotyls, reduced amounts of dephosphorylated BZR1, and decreased enrichment of BZR1 at known BZR1-target genes (fig. S5, A to D), confirming the effective blocking of the BR signaling by PPZ treatment.

We then carried out ChIP-seq using 5-day-old *BRM-GFP* transgenic seedlings treated with dimethyl sulfoxide (DMSO) or PPZ in the dark. PPZ treatment resulted in a significant decrease in BRM binding near the TSSs of a set of genes (top 50 and top 100) showing decreased chromatin accessibility in *bri1-701* mutants (Fig. 4, A and C). The enrichment of BRM also performed a significant decrease at all *bri1-701* decreased accessibility genes (fig. S6, A and B). In contrast, we did not observe significant changes in BRM binding in PPZ-treated plants at genes with increased chromatin accessibility in *bri1-701* mutants (Fig. 4, D to F and fig. S6, A and B). When we repeated the analysis using enrichment relative to binding peaks, rather than TSSs, we observed a significant decrease in BRM binding at *bri1-701* decreased accessibility genes (top 50, top 100, and all), but no significant changes in BRM binding at *bri1-701* increased accessibility genes (top 50, top 100, and all) (Fig. 4, D to F and fig. S6, C and D). At the single-gene level, genome browser snapshots of BRM ChIP–seq reads at the selected genes showed BR-deficiency-induced reduction of BRM binding at *bri1-701* decreased but not increased accessibility genes (Fig. 4G), and these results were independently validated by ChIP-qPCR analysis (Fig. 4, H and I). Notably, the BRM-GFP mRNA and protein levels were not significantly altered after PPZ treatment (Fig. 4, J and K), suggesting that the observed reduction in BRM binding at *bri1-701* decreased accessibility genes upon the loss of BR signaling was not due to the changes in BRM protein abundance. In addition, we observed a significant decline of chromatin accessibility at genomic sites showing decreased BRM binding in the absence of BR signaling; however, there was no significant changes in chromatin accessibility at genomic sites with enhanced BRM binding in the absence of BR signaling (fig. S7, A to D). These data suggest that downregulation of chromatin accessibility due to BR deficiency is directly associated with decreased BRM binding. Altogether, these results highlight the role of the BZR1-BAS complex interaction in directing BAS complex localization and remodeling activities to the *bri1-701* decreased chromatin accessibility sites.

**Fig. 4.**
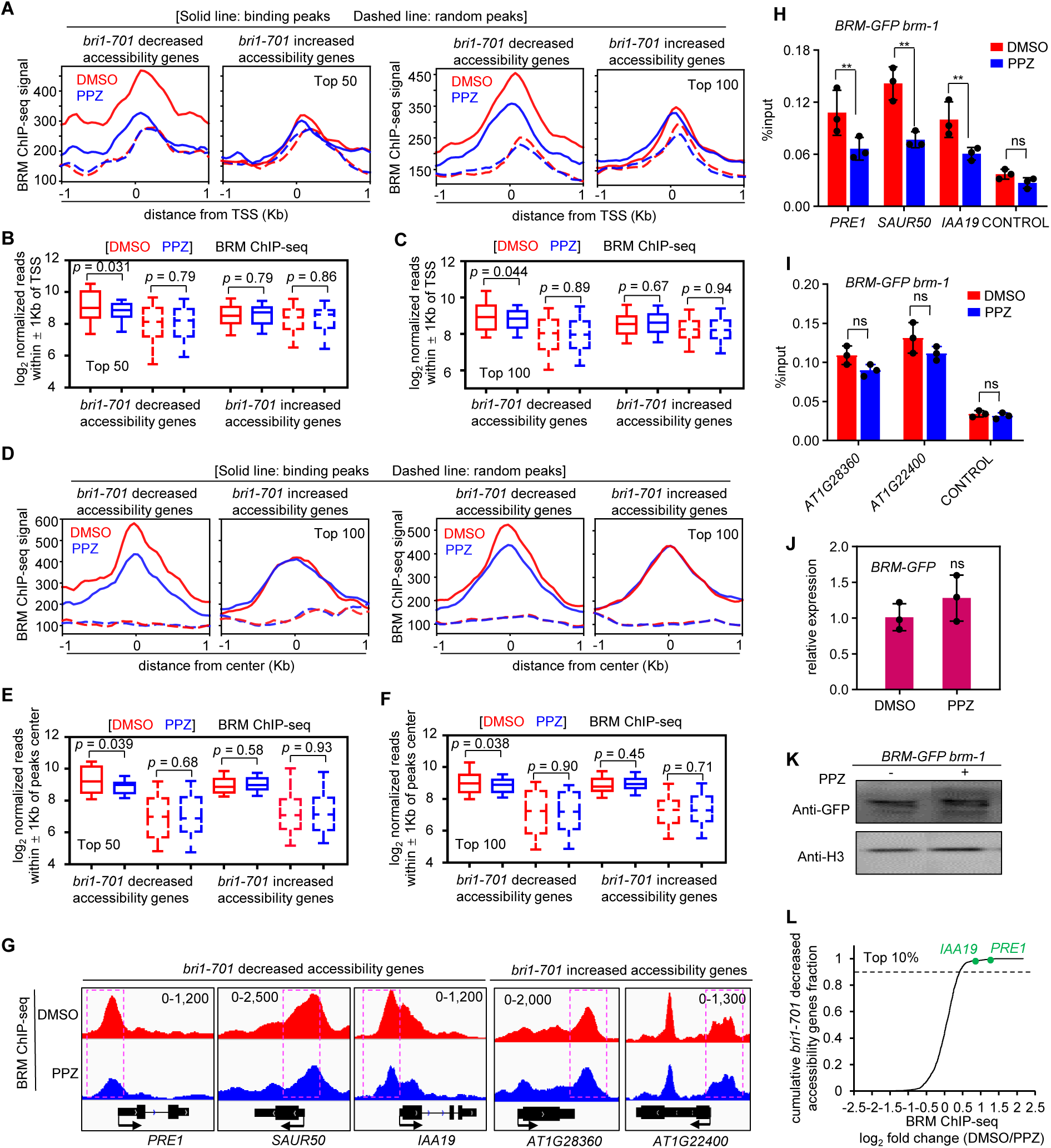
Enhanced BRM targeting is mediated by BR at *bri1-701* decreased accessibility sites. (**A**) Metagene plots displaying the ChIP-seq signals of BRM at the TSS of 50 genes (top 50) or 100 genes (top 100) showing decreased or increased accessibility in the *bri1-701* mutants. (**B**), (**C**) Box plots displaying read counts of BRM-ChIP-seq data for Top 50 or Top 100 bri1-701 decreased or increased accessibility genes. Reads were summed ± 1 Kb from the TSS. Significance analysis was determined by two tailed Mann-Whitney U test. (**D**) Metagene plots displaying the ChIP-seq signals of BRM binding peaks at 50 genes (top 50) or 100 genes (top 100) showing decreased or increased accessibility in the *bri1-701* mutants. (**E**), (**F**) Box plots displaying read counts of the BRM ChIP-seq data at top 50 or top 100 *bri1-701* decreased or increased accessibility genes. Reads were summed ± 1 Kb from the peak center. Significance analysis was determined by two tailed Mann-Whitney U test. (**G**) IGV view of ChIP– seq reads of BRM at the *bri1-701* decreased or increased accessibility genes. The black diagrams underneath indicate gene structure. The y-axis scales represent shifted merged MACS2 tag counts for every 10-bp window. (**H**), (**I**) Validation of the occupancy at the selected sites by ChIP-qPCR in the indicated transgenic plants. Mean ± s.d. from three biological replicates. Statistical significance was determined by two-tailed Student’s t-test; ** *p* < 0.01. ns, not significant. (**J**), (**K**) RT-qPCR and immunoblot analysis showing the relative RNA and protein levels of BRM with treatment of DMSO or 2µM PPZ. (**L**) Cumulative distribution function plot reflecting genes nearest to BRM decreased sites in *bri1-701* decreased accessibility genes, the top one-tenth fraction reflects genes associated with the top changed genes.

Finally, we wanted to identify sites showing most highly decreased BRM targeting and chromatin accessibility within *bri1-701* decreased accessibility genes as a strategy to identify gene loci that may underpin the biological relevance of BAS complexes in BR signaling pathway. We ranked the decreased BRM binding sites upon treatment of PPZ versus DMSO treatment and identified genes closest to these sites. This strategy led us to identify the top-regulated loci including *PRE1* and *IAA19* (Fig. 4L), which were reported to mediate cell elongation processes, thus suggesting that BZR1-BAS complex may regulate a cell elongation gene expression axis, a well-known function of BR signaling pathway.

### Disruption of BAS complex activity blocks BR-mediated chromatin accessibility enhancement

To investigate the essentiality of BRM in BR-mediated regulation of chromatin accessibility, we analyzed the impact of the loss of BRM on DNA accessibility in BR regulated chromatin accessibility regions. The chromatin accessibility levels at *bri1-701* decreased accessibility peaks were also significantly reduced in *brm-1* mutants (fig. S8, A and C). In contrast, no significant increase was observed in *brm-1* mutants at *bri1-701* increased accessibility peaks (fig. S8, B and C). Genome browser snapshots of ATAC–seq reads exemplified these results at the single-gene level (fig. S8D). Hence, these results imply that BRM may play a role in mediating BR-signaling-driven chromatin accessibility increase rather than decrease.

We next determined whether BR-mediated changes in chromatin accessibility requires BRM. We performed ATAC-seq using Col, *bzr1-1D*, *brm-1*, *bzr1-1D brm-1* seedlings grown on the medium containing 2 μM PPZ for five days in the dark. *bzr1-1D* is a gain-of-function mutant BZR1 protein that harbors a proline at position 234 to leucine substitution (P234L) (*50*), which causes BZR1 stabilization and accumulation in the nucleus (*9*). We identified a cluster of 2,494 sites over which accessibility increased in *bzr1-1D*, along with another cluster of reduced sites (n = 844) (fig. S9, A and B and table S1). Most of these differential peaks were also located upstream or downstream of genes, consistent with those peaks in *bri1-701* mutants (fig. S9C). Notably, sites showing increased and reduced accessibility in *bzr1-1D* exhibited reduced and increased accessibility, respectively, in the *bri1-701* mutants (fig. S9, D to G).

Compared with *bzr1-1D* in WT background, *bzr1-1D* nearly lost the ability to enhance chromatin accessibility in *brm-1* background, because the upregulation of accessibility by *bzr1-1D* was abolished to a Col level in the absence of BRM (Fig. 5, A and B). Heatmap analysis confirmed that disruption of BRM completely blocked the ability of *bzr1-1D* to increase chromatin accessibility (Fig. 5C). These results demonstrate that the ability of BR to increase chromatin accessibility is entirely dependent on BRM. On the contrary, when we analyzed *bzr1-1D*-reduced accessibility sites, we found that *bzr1-1D* in *brm-1* background still significantly reduced the chromatin accessibility at these sites, as it did in Col background (Fig. 5, D and E). Heatmap analysis again showed that the loss of BRM largely did not disturb the ability of bzr1-1D to decrease chromatin accessibility (Fig. 5F). These results support the notion that BRM activity is largely not required for the ability of BR to decrease chromatin accessibility. PCA analysis showed that *bzr1-1D brm-1* grouped with Col at BZR1 increased chromatin accessibility sites, while, at BZR1decreased chromatin accessibility sites, *bzr1-1D brm-1* was more associated with *bzr1-1D* (Fig. 5, G and H). At the single-gene level, genome browser snapshots of ATAC-seq reads confirmed that BRM is required for BZR1 to increase chromatin accessibility at *PRE1* and *IAA19* genes but not to decrease chromatin accessibility (Fig. 5I). Taken together, these data demonstrate that BRM is essential for increasing rather than decreasing chromatin accessibility of genes regulated by the BR signaling pathway.

**Fig. 5.**
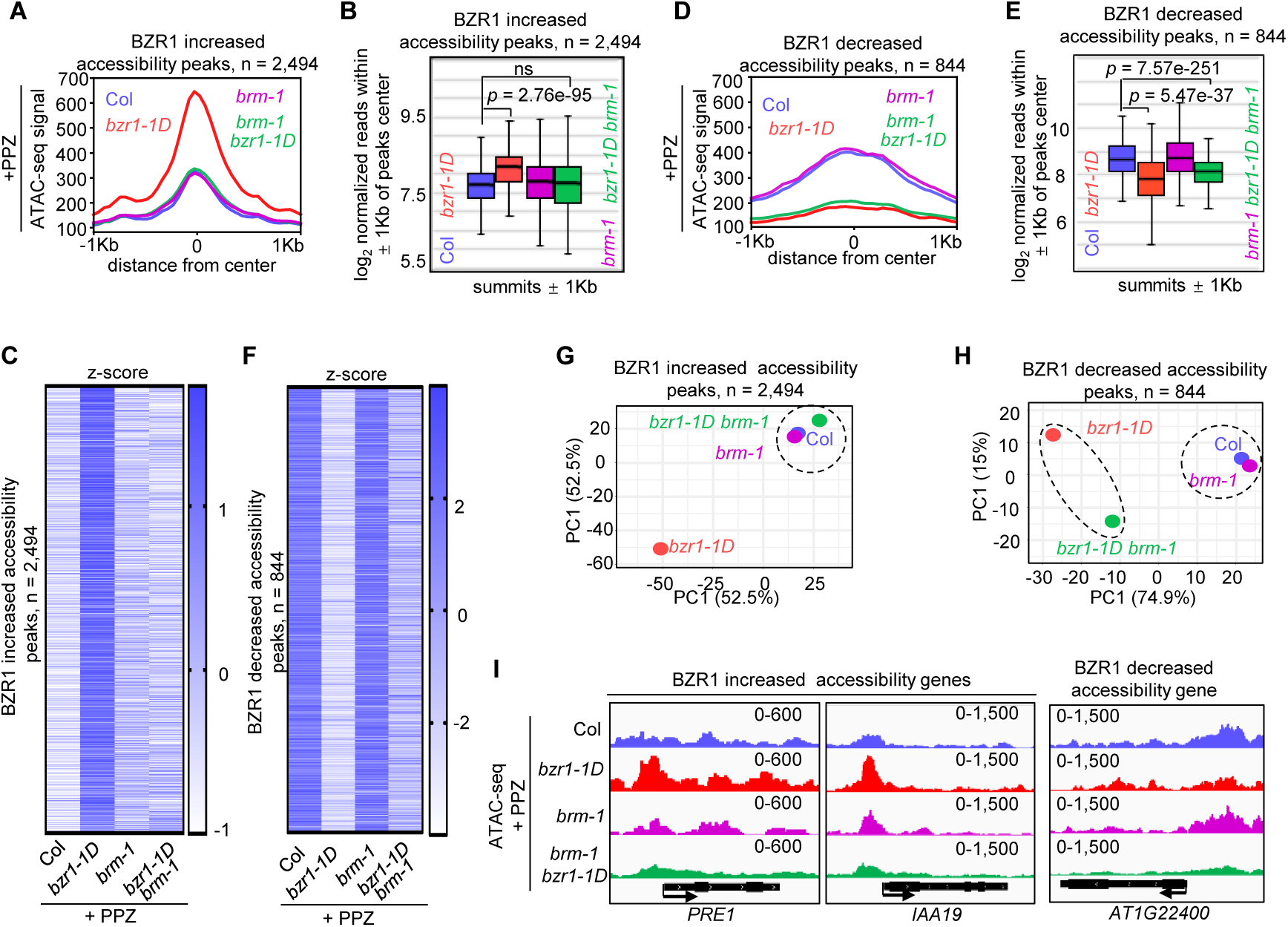
BZR1 requires BRM to increase chromatin accessibility. (**A**) Metagene plots reflecting the ATAC-seq signals over BZR1 increased chromatin accessibility peaks for the indicated ATAC-seq experiments. Seedlings were grown in the dark with 2µM PPZ for five days. (**B**) Box plot displaying read counts over the BZR1 increased chromatin accessibility peaks for the indicated ATAC-seq experiments. Reads were summed ± 1 Kb from the peaks center. Significance analysis was determined by two tailed Mann-Whitney U test. (**C**) Heatmap reflecting the ATAC-seq signals over the increased chromatin accessibility peaks by *bzr1-D* for the indicated ATAC-seq experiments. (**D**) Metagene plots reflecting the ATAC-seq signals over BZR1 decreased chromatin accessibility peaks for the indicated ATAC-seq experiments. Seedlings were grown in the dark with 2µM PPZ for five days. (**E**) Box plot displaying read counts over the BZR1 decreased chromatin accessibility peaks for the indicated ATAC-seq experiments. Reads were summed ± 1 Kb from the peaks center. Significance analysis was determined by two tailed Mann-Whitney U test. (**F**) Heatmap reflecting the ATAC-seq signals over the decreased chromatin accessibility peaks by *bzr1-D* for the indicated ATAC-seq experiments. (**G**), (**H**) PCA analysis of bzr1-1D increased or decreased accessibility peaks in Col, *bzr1-1D*, *brm-1* and *bzr1-1D brm-1* samples. Percentages represent variance captured by PC1 and PC2 in each analysis. (**I**) Examples of ATAC-seq tracks at representative loci in the Col, *bzr1-1D*, *brm-1* and *bzr1-1D brm-1* samples.

### BRM deficiency blocks the elongation of hypocotyl and downregulates the expression of cell elongation-related genes

We next sought to define the biological function underpinning the essentiality of the BZR1-BAS complexes in BR-mediated chromatin accessibility. Similar to the *bri1-701* mutants, the *brm-1* mutants displayed a strongly reduced hypocotyl length under dark conditions (fig. S10A). Inactivation of the BAS-specific subunits (*brip1 brip2* double or *brd1 brd2 brd13* triple mutants) also impaired hypocotyl elongation under dark conditions (fig. S10B). These results implied that BAS complexes may be required for the BR-mediated hypocotyl elongation. Indeed, the *brm-1* mutants were more sensitive to PPZ treatment (Fig. 6A), suggesting that loss of BRM compromises BR responses.

**Fig. 6.**
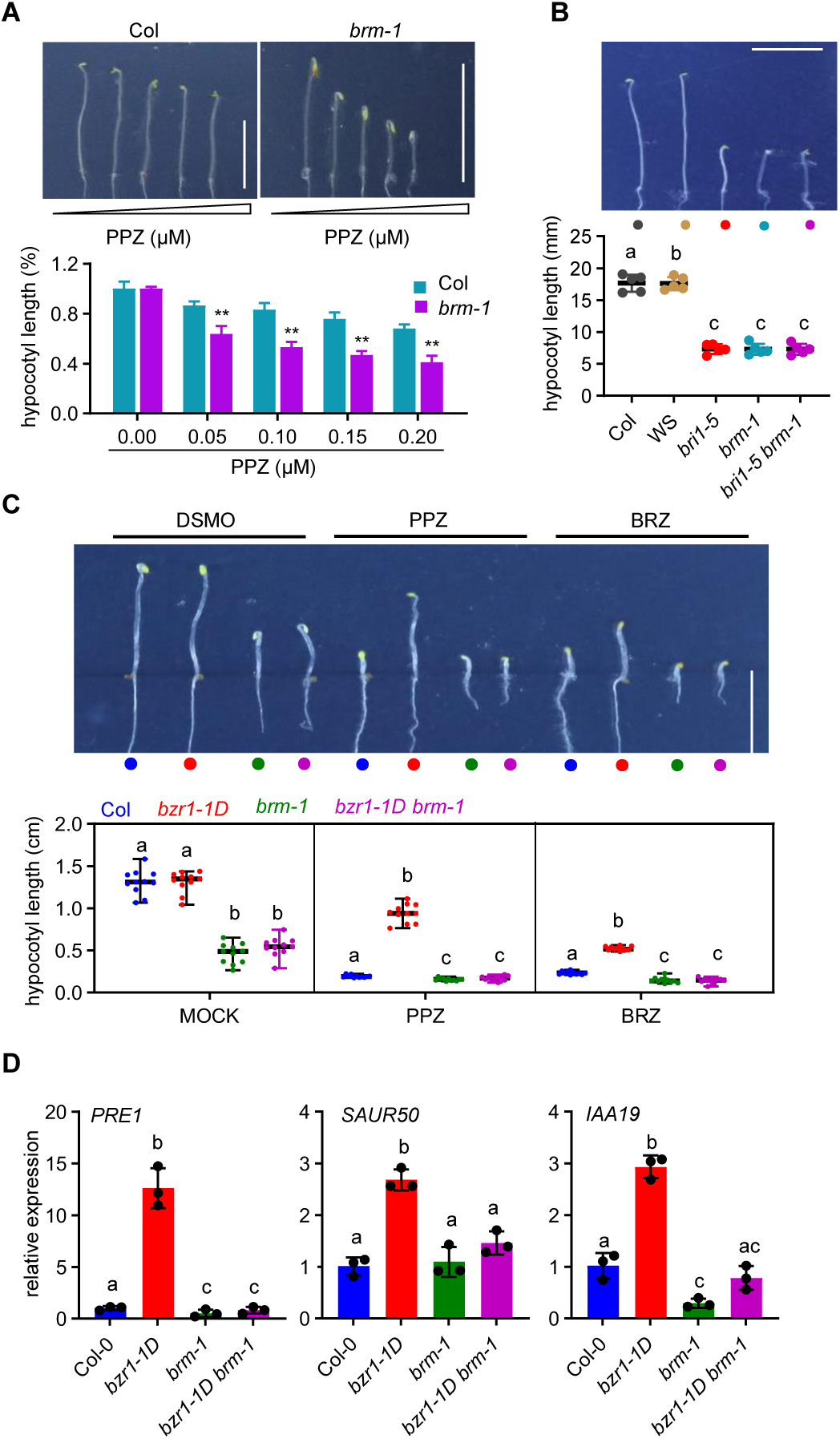
Loss of BRM compromises the elongation of hypocotyl and downregulates cell elongation related genes. (**A**) The *brm-1* mutant is hypersensitive to PPZ. Seedlings were grown on various concentrations of PPZ in the dark for five days. The error bars in the lower graph indicate the s.d. (n = 10 plants) and \*\**p* < 0.01. Scale bar, 10 mm. (**B**) The loss of BRM inhibits the promotion of hypocotyl elongation. Seedlings were grown in the dark for five days. The error bars in the lower graph indicate the s.d. (n = 10 plants). Lowercase letters indicate statistical significance determined by the Student’s t test. Scale bar, 10 mm. (**C**) *bzr1-1D* under the brm-1 background cannot promote hypocotyl elongation in the dark. Seedlings were grown on medium either with DMSO or 2 µM PPZ or 2 µM BRZ in the dark for five days. The error bars in the lower graph indicate the s.d. (n = 10 plants). Scale bar, 10 mm. (**D**) Relative expression of *PRE1*, *SAUR50* and *IAA19* in 5-day-old seedlings grown in dark conditions. *ACTIN2* served as the internal control. Mean ± s.d. from three biological replicates. Lowercase letters indicate statistical significance determined by the Student’s t test.

We further generated *bri1-5 brm-1* double mutants to investigate the genetic relationship between BZR1 and BRM. Comparison of hypocotyl length of the double mutant *bri1-5 brm-1* with that of the *bri1-5* and *brm-1* single mutants showed that loss of the BR receptor BRI did not exacerbate the shortened hypocotyl phenotype of the *brm-1* mutants (Fig. 6B), suggesting a role for the BAS complexes operated through the BR-signaling pathway to regulate hypocotyl elongation in the dark. In support of this notion, the *bzr1-1D brm-1* double mutants had hypocotyl length similar to *brm-1* grown on the medium with or without PPZ or BRZ (*51*) (a specific BR biosynthesis inhibitor—brassinazole) (Fig. 6C), demonstrating that the BZR1-BAS interaction is part of the BR-signaling pathway in regulating hypocotyl elongation in the dark.

We carried out reverse transcription followed by quantitative PCR (RT-qPCR) of several cell-elongation-associated genes in Col, *bzr1-1D*, *brm-1*, *bzr1-1D brm-1* mutants to explore the role of BRM in regulating the expression of cell-elongation associated genes. As shown in Fig. 6D, disruption of BRM severely compromised the *bzr1-1D*-induced upregulation of cell-elongation-associated genes, including *SAUR50*, *IAA19*, and *PRE1*. Taken together, these data indicate that the loss of BRM blocks the BZR1-promotion of hypocotyl elongation and downregulates the expression of genes involved in cell elongation.

### BRM is required for the expression of BR-activated, but not BR-repressed, genes

To further clarify the genome-wide role of the BZR1-BAS complexes in BR-regulated transcriptional activation or repression processes, we conducted RNA-sequencing (RNA-seq) assay using Col, *bzr1-1D*, *brm-1*, *bzr1-1D brm-1* seedlings grown on the medium containing 2 μM PPZ in the dark for five days. We identified 929 upregulated and 1,096 downregulated genes (|log_2_ fold change| ≥ 1) affected more than twofold by *bzr1-1D* mutation when compared with Col (Fig. 7A and table S3). Profiling the dependence of the genes upregulated by bzr1-1D on BRM allowed us to define three clusters of genes including: those activated by BZR1 and repressed by BRM for expression (cluster 1, 19%); those moderately dependent on BRM for BZR1-mediated activation (cluster 2, 31%); and those entirely depends on BRM for BZR1-mediated activation (cluster 3, 50%) (Fig. 7B). Therefore, most of BZR1-upregulated genes (81%,753 out of 929) no longer or less upregulated in the *brm-1* background (Fig. 7, B and D), suggesting the critical role for BRM in mediating the transcriptional activation of BZR1. However, when we analyzed the genes repressed by *bzr1-1D* in the Col background, we found the majority of them (855 out of 1096, 78%) were still down regulated in the *brm-1* background (Fig. 7, C and D), suggesting that BZR1-mediated transcriptional repression is largely independent of BRM.

**Fig. 7.**
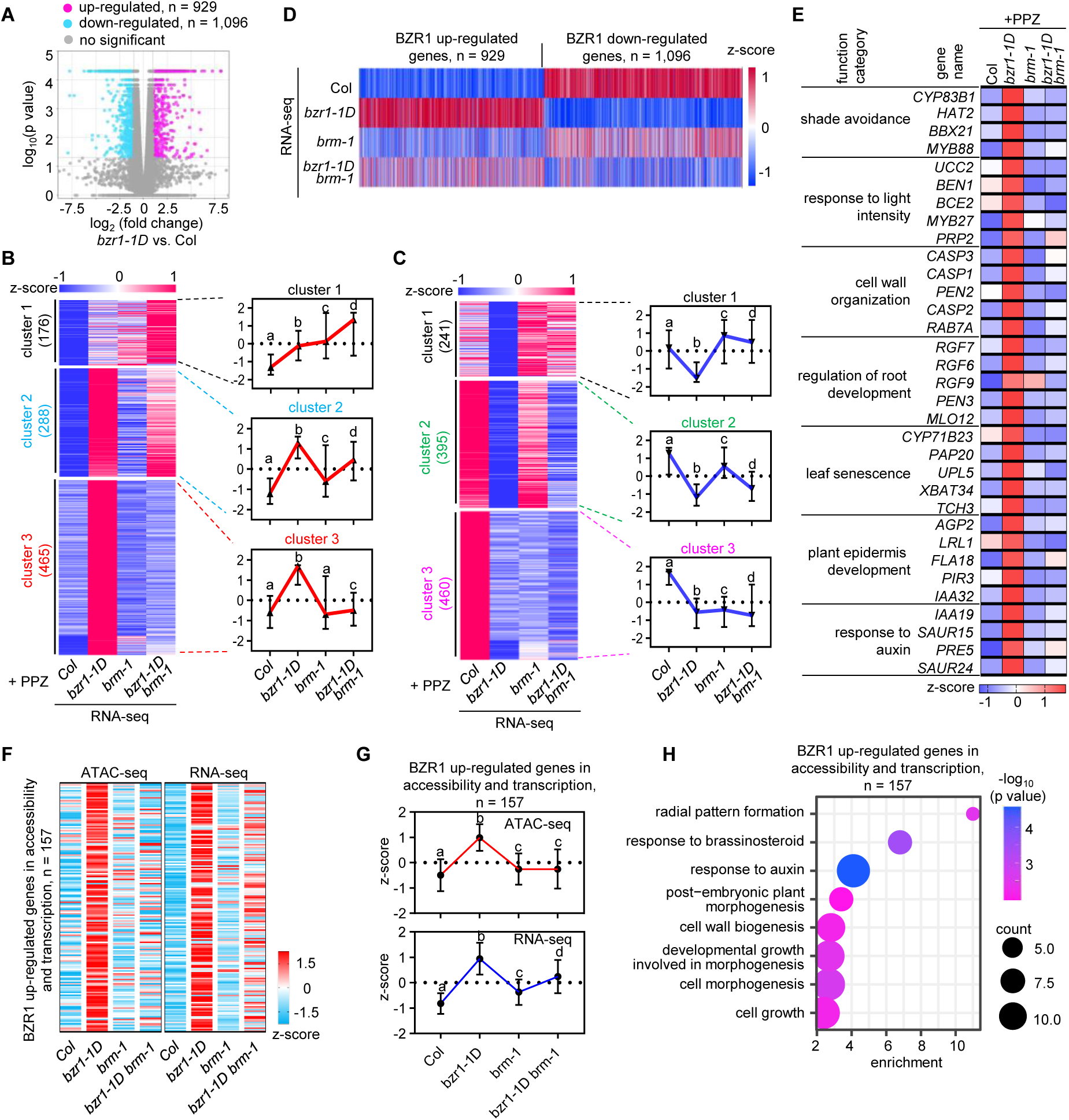
BRM determines BR-mediated gene activation, but has no effect on BR-mediated gene repression to a large extent. (**A**) Volcano plots showing differentially expressed genes in the *bzr1-1D* mutants, determined by RNA-seq. |log_2_(fold change)| ≥ 1. (**B**) Heatmap (left) and box blot (right) showing the classes of BZR1 up-regulated genes sorted by k-means clustering across the samples collected from Col, *bzr1-1D*, *brm-1*, *bzr1-1D brm-1* samples. Color bar, RNA z-score of the differentially expressed genes identified by RNA-seq. The number of the genes for each cluster is given. (**C**) Heatmap (left) and box blot (right) showing the classes of BZR1 down-regulated genes sorted by k-means clustering across the samples collected from Col, *bzr1-1D*, *brm-1*, *bzr1-1D brm-1* samples. Color bar, RNA z-score of the differentially expressed genes identified by RNA-seq. The number of the genes for each cluster is given. (**D**) Heatmaps reflecting the relative expression (*z*-normalized) of BZR1 up or down-regulated genes for the indicated RNA-seq experiments. (**E**) Comparative expression analyses of BRM-dependent BZR1 up-regulated genes in diverse developmental programmers. Heatmap of RNA-seq data from triplicate biological samples prepared from Col, *bzr1-1D*, *brm-1*, *bzr1-1D brm-1* seedings. (**F**) Heat map displaying the chromatin accessible and transcriptional relative level (*z*-normalized) at BZR1 up-regulated genes in both transcription and chromatin accessibility for indicated samples. z-score values of chromatin accessibility and gene expression in indicated samples were also displayed. (**G**) Box blot displaying the chromatin accessible and transcriptional relative level (*z*-normalized) at BZR1 up-regulated genes in both transcription and chromatin accessibility for indicated samples. (**H**) Gene ontology analysis using 157 genes in **F**.

GO analysis of the 753 BRM-dependent BZR1-upregulated genes revealed significant enrichment controlling a broad spectrum of developmental programmers including shade avoidance, root development, leaf senescence, response to light intensity, cell wall organization, epidermis development and response to Auxin, all of which are well-known BR-regulated pathways (Fig. 7E). In particular, Upregulation of genes involved in shade avoidance, light response, and cell wall organization in *bzr1-1D* was diminished in the *bzr1-1D brm-1* mutants. The aberrant upregulation of genes related to root development in *bzr1-1D* was abolished by the disruption of BRM. We also observed the activated expression of leaf senescence and plant epidermal development-related genes by BZR1 depends on BRM. Important genes governing auxin responses, including *IAA19*, *SAUR50*, and *PRE1*, were unable to be activated by BZR1 upon the loss of BRM.

Furthermore, integration of ATAC-seq and RNA-seq datasets identified a subcluster of genes (*n* = 157) that showed up-regulated transcription and DNA accessibility in *bzr1-1D* mutants (Fig. 7F). The increase in RNA expression and accessibility by bzr1-1D was largely dependent on BRM (Fig. 7F). GO term analysis of these genes also revealed terms related to growth and development processes, such as radial pattern formation, response to Auxin and BR, cell wall organization, epidermis development and so on (Fig. 7G). Together, these data demonstrate that BZR1-BAS complexes have a vital role in gating BR-responsive genome accessibility and transcriptional activation in diverse post-embryonic developmental programs throughout plant life.

## Discussion

As a master transcription factor in the BR signaling pathway, BZR1 regulates the expression of thousands of genes involved in diverse developmental and stress response programs. However, in contrast to the well-characterized BR signaling pathway upstream of BZR1, the downstream mechanisms by which BZR1 regulates gene expression are less understood. In this study, we report a direct molecular connection between BR-BZR1 signaling and SWI/SNF regulation (Fig. 8). BR signaling can modulate the chromatin accessibility and the consequential activation or repression of transcription of thousands of genes regulated by BZR1 (Fig. 1 and fig. S1). Mechanistically, we show that nucleus-localized BZR1 physically interacts with BRM and several BAS-specific subunits (Fig. 2 and fig. S2), has a high colocalization with BRM on the genome (Fig. 3 and fig. S3), and enhances BRM occupancy at BR increased accessibility sites (Fig. 4 and fig. S6). BRM governs the BR-mediated chromatin accessibility increase, rather than decrease (Fig. 5). Finally, phenotypic and transcriptome analysis provided compelling evidence for the indispensability of BRM in BR-mediated hypocotyl elongation (Fig. 6) and genome-wide transcriptional activation (Fig. 7). We propose that BAS-SWI/SNF chromatin remodeling acts to dictate the transcriptional activation activity of BZR1 for BR-regulated growth and development in plants. This signaling axis thus serves as a phytohormone-mediated checkpoint for regulating BAS-SWI/SNF activity essential for key developmental phases and processes throughout plant growth and development (Fig. 8A).

**Fig. 8.**
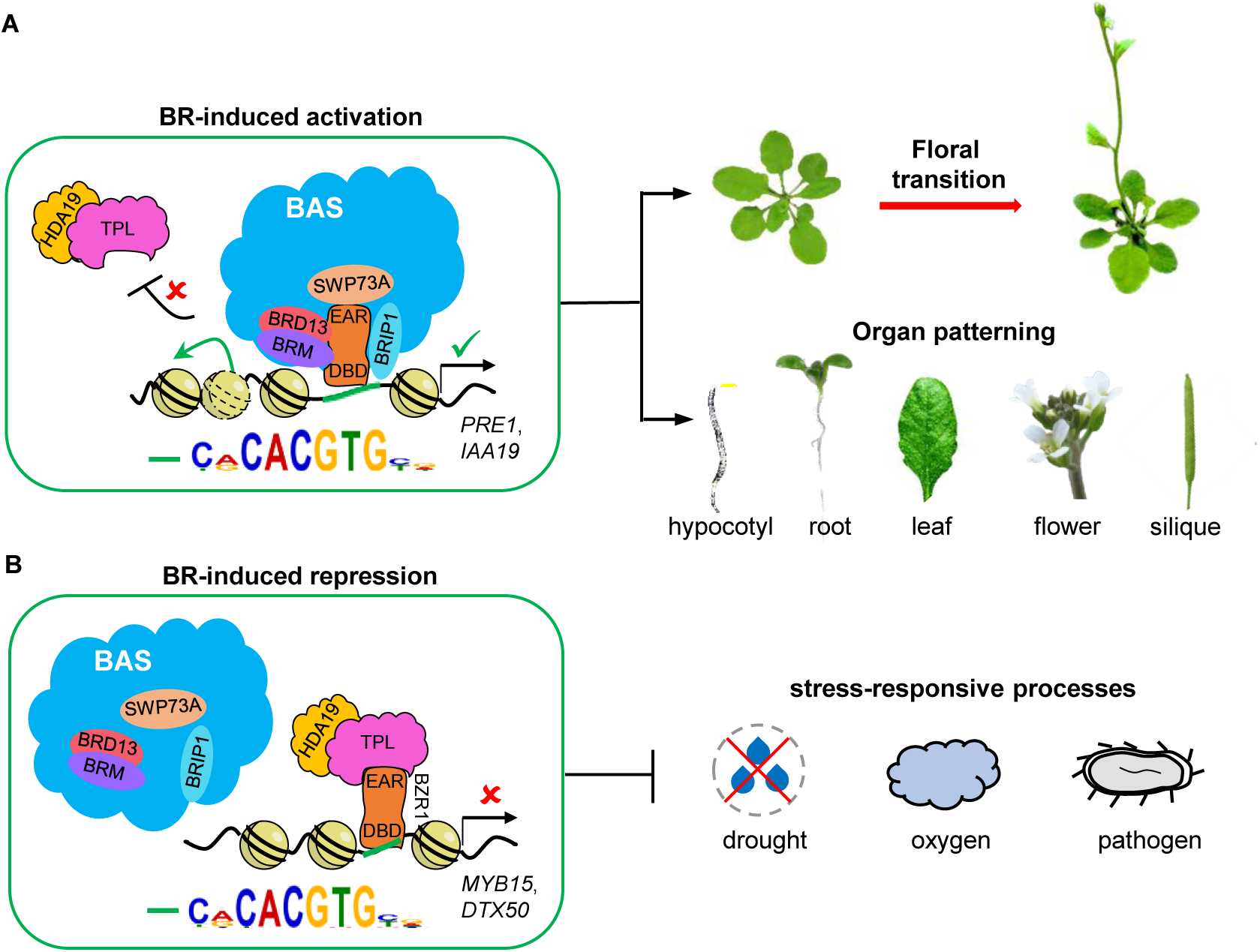
Model of the BR-BZR1-BAS signaling network governing diverse developmental programs. (**A**) The BR-BZR1-BAS-mediated transcriptional activation signaling network. BR activated BZR1 interacts with and recruits the BAS complexes to the G-box-like containing genes, where BAS enhances chromatin accessibility and activate gene expression to support a range of plant growth and developmental processes, including fruit and seed development, hypocotyl elongation, root growth, leaf expansion, flowering transition, and floral organ formation. This molecular mechanism establishes a direct and global mechanistic connection between hormones and chromatin accessibility during plant growth and development process. (**B**) The BR-BZR1-TPL-HDA19-mediated transcriptional repression signaling network. BZR1-TPL-HDA19 complexes bind to the G-box-like motifs in the stress-responsive genes and inhibit their expression to help balance the trade-off between growth and stress response.

Plants are constantly exposed to various stress signals in facing their environment, and thus, they must balance their growth and defense mechanisms to optimize fitness (*52*). Understanding the balance mechanism of growth and defense is important for developing strategies to maximize crop yield (*53, 54*). BR has been identified as a critical hormone in plant growth-defense coordination; however, the underlying mechanisms remain poorly understood (*10, 55–57*). Our genome-wide analyses define a broad spectrum of development-related pathways as targets of the BZR1-SWI/SNF signaling network (fig. S1J and Fig.7E). This BR-stimulated epigenomic activation network induces transcription of genes required for promoting growth-related programs, including shade avoidance, root development, cell wall organization, epidermis development, and response to light intensity and Auxin (Fig.7E). By contrast, pathways associated with responses to hypoxia, salicylic acid, salt stress, and oxidative stress are the major targets of BR-mediated chromatin repression (fig. S1K). This dual molecular role of BR in balancing chromatin accessibility states between growth- and stress related genes ensures that the proliferation and differentiation processes during plant growth in time and space are coordinated and balanced with stress conditions (Fig. 8, A and B).

Notably, recent studies reported that hormones other than BR are also associated with chromatin accessibility regulation in plants. Cytokinins (CK) regulate the development of specific plant tissues by modulating chromatin accessibility (*58*). Auxin plays a pivotal role in triggering dynamic changes in chromatin accessibility during embryonic development (*59*). However, the precise molecular mechanisms governing these hormone-induced alterations in chromatin accessibility remain enigmatic. Given the reported functional connection between CK or Auxin and BRM remodelers (*60, 61*), it is tempting to speculate that a parallel mechanism involving SWI/SNF-mediated chromatin remodeling activity may be responsible for these phytohormones to govern epigenetic landscapes and gene regulatory dynamics in plants.

What determines the transcriptional activation versus repressive activity of BZR1 is a long-standing question. Previous in vitro DAP-seq analysis suggested that the motifs recognized by BZR1 might determine its transcriptional activity, with BZR1 preferentially recognizes the 10 bp DNA fragment containing the known G-box core binding motif at the center for transcriptional repression activity; however, the mechanism of BZR1-induced transcriptional activation is unknown (*29*). Here, we found through integrating ChIP-seq and ATAC-seq data that BZR1 significantly enriches in both BR-decreased and increased DARs (Fig. 1I). Further analysis revealed that a same 10 bp DNA sequence, containing a G-box core motif, was significantly enriched in both the increased and decreased DARs (Fig. 1, F and G), implying that *cis* motif may not be the major determinant of the transcriptional activation or repression activity of BZR1. Therefore, other factors are assumed to be responsible for distinguishing the mutual transition between transcriptional activation and repression of BZR1. Here, we found that BRM mediates the transcriptional activation ability of BZR1 through increasing chromatin accessibility but does not involve in the transcriptional repression of BZR1, indicating that BRM as a transcriptional co regulator that specifically confers the transcriptional activation activity of BZR1. Interestingly, previous studies have shown that TPL, acting as a co-repressor, interacts with the EAR domain of BZR1 to determine its transcriptional repression activity (*30*). Therefore, we propose that trans-regulators rather than cis-elements determine BZR1’s transcriptional activation and repression activity.

Interestingly, the EAR domain of BZR1 is required not only for transcriptional repression but also for transcriptional activation (*30, 62*). However, the mechanistic action of EAR in BZR1 transcriptional activation is unknown. Surprisingly, although our results showed that BRM ATPase of the BAS complex interacts with the N-terminal region of BZR1, the core subunits BRIP1, BRD13, and SWP73A of the BAS interact with the EAR domain-containing C-terminal region of BZR1 (Fig. 2G). Moreover, BAS interacts with the EAR domain of BZR1 through the SWP73A core subunit (fig. S2H). Thus, given that both the BAS subunits and TPL can interact with the EAR domain, we propose that when BZR1-BAS activates genes, the interaction between BAS and BZR1 on the promoter of BR-activated genes may trigger a conformational change that prevents the EAR domain from being approached by the TPL co-repressor. This transition of transcriptional co-regulators, from TPL to BRM, may determine the transcriptional activation capacity of the EAR domain. Given the ubiquity of the EAR motif in multiple transcription factors with a dual function in activation and repression, it will be interesting to examine whether BAS may be responsible for transmitting the transcriptional repression function to the activation ability of diverse transcription factors.

Our data show that, in addition to its role in inducing chromatin accessibility, BZR1 is also able to repress DNA accessibility (Fig. 5, D and F). Loss of BRM largely does not disturb the function of BZR1 in decreasing the chromatin accessibility (Fig. 5, D and F), implying that the BZR1-mediated decrease in chromatin accessibility likely requires other chromatin regulators. Apart from BAS, Arabidopsis has two other subcomplexes of the SWI/SNF complexes, SAS and MAS. However, like BAS, the SAS and MAS are primarily involved in enhancing chromatin accessibility (*41*), therefore, are unlikely to be responsible for the BR-mediated DNA accessibility downregulation. Other candidates could be imitation switch (ISWI), chromodomain helicase DNA-binding (CHD), and inositol requiring 80 (INO80) remodeling complexes, although whether they can regulate chromatin accessibility in the genome remains unclear. In addition, previous studies have demonstrated that TPL and HDA19 are responsible for BR-mediated transcriptional repression. Thus, the possibility that BZR1 may rely on the TPL-HDA19 module to confer a closed chromatin landscape requires further evaluation.

In summary, our work uncovers the mechanistic basis for the transcriptional activation activity of the BR signaling pathway. This molecular mechanism provides a long-sought mechanistic explanation for how BR signaling activates multiple developmental processes including hypocotyl elongation in plants. Our study advances a conceptual understanding of how multicellular organisms convert systemic hormonal information to remodel the global chromatin accessibility landscapes by modulating local chromatin regulators, thus orchestrating transcriptional states that are central for diverse developmental programs.

## Materials and Methods

### Plant materials and growth conditions

The mutants *brm-1* (SALK_030046), *brip1 brip2* (SALK_133464 and SALK_177513), *brd1 brd2 brd13* (SALK_1012963, SALK_025965 and SALK_053556), and *pBRM:BRM-GFP brm-1* transgenic plants were previously described (*47, 48, 63*). The *pBZR1:BZR1-YFP* transgenic plants were previously described (*15*). The *bri1-5* (*64*) mutants were kindly provided by Prof. Hongwei Xue. The *bzr1-1D* (*50*) mutants were kindly provided by Prof. Junxian He. The *bri1-701* mutants were kindly provided by Prof. Jia Li. All plants were in the Columbia-0 (Col-0) background except for *bri1-5*, which is in the Wassilewskija (Ws) ecotypes.

Arabidopsis plants were grown in a greenhouse with a 16-h light/8-h dark cycle at 22 °C for general growth and seed harvesting. For RT-qPCR/RNA-seq, ChIP-qPCR/ChIP-seq, ATAC-seq, and IP-MS assays, seeds were subjected to a sterilization process using a 15% sodium hypochlorite solution, followed by three washes with sterile water. Subsequently, the sterilized seeds were stratified in darkness at 4 °C for a duration of three days. The stratified seeds were then sown onto ½-strength Murashige and Skoog (MS) medium supplemented with 1% sucrose and 0.6% agar.

### Generation of transgenic plants

For *BZR1-3xFLAG BRM-GFP brm-1*, genomic regions corresponding to full-length BZR1, including a 2.0-kb promoter and the coding region without the stop codon, were amplified and subcloned into *pZPY122-FLAG* (*65*) (after cutting with restriction enzymes KpnI and PstI) using a homologous recombination with the ClonExpress Entry One Step Cloning Kit (Vazyme, Cat. No. C114). The construct was introduced into A. tumefaciens strain *GV3101*, which was used to transform *pBRM:BRM-GFP brm-1* transgenic plants using the floral dip method. Primers used for constructing are listed in table S4.

### Hypocotyl length measurements

Seeds were subjected to sterilization using a 15% sodium hypochlorite solution, followed by cultivation on half-strength MS medium supplemented with 0.8% agar. After a cold stratification period of three days at 4 °C, the seedlings were exposed to white light for 6 h to induce germination. Subsequently, the seedlings were incubated under dark condition for five days. Photocopies of the seedlings were obtained, and the lengths of the hypocotyls were measured using ImageJ software (http://rsb.info.nih.gov/ij).

### Y2H assays

The BAS-associated Y2H vectors employed in this study have been previously described (*47, 48*). To generate full-length and truncated versions of BZR1, the corresponding truncated fragments were amplified from Col cDNA and subsequently cloned into the BamHI sites of *pGADT7* or *pGBKT7* plasmids using the ClonExpress II One Step Cloning Kit (Vazyme, Cat. No. C112-01). The resulting constructs were co transformed into the Y2H Gold yeast strain (*AH109*), and all yeast cells were cultured on selective media, such as SD medium lacking leucine and tryptophan or SD medium lacking adenine, histidine, leucine, and tryptophan. Primers used for constructing are listed in table S4.

### BiFC assays

The full-length coding sequences of BZR1 were amplified from cDNA derived from Arabidopsis thaliana Col-0 and subsequently inserted into the *pEarleyGate 201-nYFP* vector (*66*) using the LR reaction (Invitrogen). The BAS-associated BiFC vectors employed in this study have been previously described (*47, 48*). The resulting constructs were individually introduced into Agrobacterium tumefaciens strain *GV3101*, and the transformed bacteria were then used for infiltration into the lower epidermal cells of Nicotiana benthamiana leaves (*67*). After a 48-hour incubation period, YFP fluorescence signals were visualized using a confocal microscope (LSM880 with Fast Airy scan). As a negative control, HAT3 (encoded by *AT3G60390*) was included. Primers used for constructing are listed in table S4.

### Co-immunoprecipitation

The full-length coding sequences of BZR1 and SWP73A were amplified from cDNA obtained from Arabidopsis thaliana Col-0 and subsequently inserted into the BamHI sites of the *pHB-HA* or *pHB-FLAG* vector (*68*). The vectors *pEAQ-BRM-N-GFP*, *pHB-BRIP1-HA*, *pHB-BRIP2-HA*, *pHB-BRD2-HA*, and *pHB-BRD13-HA* have been previously described (*47, 48*). Primers used for constructing are listed in table S4.

For Co-IP, the constructs were co-transformed into tobacco leaves, which were collected after 48 h. Total proteins were isolated from 0.2 g tobacco leaves and then lysed with 2 ml of IP buffer (50 mM HEPES (pH 7.5) 150 mM NaCl, 10 mM EDTA, 1% Triton X-100, 10% glycerol, 0.2% NP-40, and 1× Complete protease inhibitor cocktail (Roche)) at 4℃ for 30 min. After centrifugation at 5000 g and 4℃ for 10 min, the supernatant was incubated with 10 μl of anti-HA-agarose antibody (Sigma, Cat. No. A2095-1ML) or anti-FLAG beads (Bimake, Cat. No. B26101-1ML) at 4℃ for 3 h and then washed three times with washing buffer (50 mM HEPES (pH 7.5) 100 mM NaCl, 10 mM EDTA, 10% glycerol, 0.1% NP-40, and 1× Complete protease inhibitor cocktail (Roche)). Finally, proteins were diluted in 5× SDS loading buffer and boiled at 55℃ for 10 min, followed by immunoblotting.

For co-IP analysis of stable Arabidopsis transgenic plants, 2 g of 14-day-old seedlings grown under long-day conditions were carefully ground to a fine powder in liquid nitrogen. The resulting powder was resuspended in 30 ml of extraction buffer 1 (comprising 0.4 M sucrose, 10 mM Tris-HCl (pH 8.0), 10 mM MgCl_2_, 5 mM β-ME, 0.1 mM PMSF, and 1× Complete protease inhibitor cocktail (Roche)). The homogenate was then passed through two layers of Miracloth to remove solid debris, followed by centrifugation at 3,000 g for 20 min at 4°C. The resulting precipitates were subsequently lysed in 10 ml of IP buffer (containing 100 mM Tris-HCl (pH 7.5), 300 mM NaCl, 2 mM EDTA, 1% Triton X-100, 10% glycerol, 1 mM PMSF, and 1× Complete protease inhibitor cocktail (Roche)) at 4°C for 30 min. After centrifugation at 14,000 g for 15 min at 4°C, the supernatant was diluted with an equal volume of dilution buffer (comprising 100 mM Tris-HCl (pH 7.5), 2 mM EDTA, 10% glycerol, 1 mM PMSF, and 1× Complete protease inhibitor cocktail (Roche)). Subsequently, the diluted supernatant was incubated with anti-FLAG beads (Bimake, Cat. No. B26101-1ML) at 4°C for 3 h with gentle rotation. The beads were then washed three times with 1×PBS solution containing 0.1% Tween-20. Finally, the proteins were eluted in 5× SDS loading buffer and incubated at 55°C for 10 min, followed by subsequent immunoblotting.

### Mass spectrometry

For mass spectrometry, the immunoprecipitated proteins using Co-IP methods were eluted using 0.2 M glycine solution (pH 2.5), and then subjected to reduction with dithiothreitol, alkylation with iodoacetamide and digested with trypsin (Thermo Fisher, Cat. No. 90057, MS748 grade). The samples were analyzed on a Thermo Scientific Q Exactive HF mass spectrometer in data-dependent mode. Spectral data were searched against the TAIR10 database using Protein Prospector 4.0. Two biological replicates were included in the IP-MS analysis. Raw data were searched against the TAIR10. Default settings for Label-free quantitation (LFQ) analysis using MaxQuant65 and Perseus software were applied to calculate the LFQ intensities with default settings.

### Immunoblotting

Protein samples were loaded onto 4%-20% gradient protein gels (GenScript, SurePAGE, Cat. No. M00655) or 10% SDS-PAGE gels and electrophoresed at 150 V for 2 h. Subsequently, a wet transfer was conducted in ice-cold transfer buffer at 90 V for 90 min. Following transfer, the membranes were blocked with 5% non-fat milk at room temperature for 1.5 h on a shaking table. The blocked membranes were then incubated with specific antibody solutions at room temperature for an additional 3 h. The antibodies used included anti-GFP (Abcam, Cat. No. ab290, diluted 1:10,000), anti-HA (Sigma-Aldrich, Cat. No. H6533, diluted 1:5000), anti-FLAG (Sigma-Aldrich, Cat. No. A8592, diluted 1:5000), anti-H3 (Proteintech, Cat. No. 17168-1-AP, diluted 1:10,000), and horseradish peroxidase-conjugated goat-anti-rabbit secondary antibody (Abcam, Cat. No. ab6721, diluted 1:10,000). The intensity of the blotting signals was quantified using ImageJ software (version 1.50i).

### RNA isolation, qRT-PCR and RNA-seq analyses

Total RNA was extracted from 5-day-old seedlings grown in the dark using the HiPure Plant RNA Mini Kit C (Cat. No. R4151-02C) following the manufacturer’s protocol. Reverse transcription reactions were carried out using 1 μg of total RNA with HiScript II Q RT SuperMix for qPCR (+gDNA wiper) with gDNA eraser (Vazyme, Cat. No. R223-01). Quantitative real-time PCR (qRT-PCR) was performed using the SYBR Green SuperMix and StepOne Software v.2.3 (Applied Biosystems) with 40 cycles of amplification (including three biological replicates). The relative expression levels were analyzed using the -ΔΔCt (cycle threshold) method (*69*), and the data were normalized to the expression of the reference gene *ACTIN2*. Primers used for constructing are listed in table S4.

For RNA-seq analyses, RNA libraries were generated using the TruSeq RNA sample preparation kit (Illumina) following the manufacturer’s instructions, and the sequencing was performed at Novogene (Beijing, China) on the Illumina novaseq PE150 platform. For data analysis, the raw sequence reads were aligned to the TAIR10 genome using TopHat (Galaxy v.2.1.1) (*70*), with a minimum intron length set to 20 and a maximum intron length set to 4000. Subsequently, the mapped reads were assembled using Cufflinks (Galaxy v.2.2.1.3) (*71*) based on the TAIR10 genome annotation, utilizing default settings. To identify differentially expressed genes, the assembled transcripts from three independent biological replicates of Col and the mutants were combined and compared using Cuffmerge (Galaxy v.2.2.1.2) (*71*) with default parameters. Genes exhibiting at least a 2-fold change in expression (false discovery rate [FDR] <0.05, P < 0.05) were considered differentially expressed and used for subsequent analysis. The heatmap and the clusters of BZR1-regulated genes sorted by k-means approach were performed using MeV software (*72*).

### ChIP experiment and ChIP–seq analysis

ChIP experiments were conducted following previously established protocols with slight modifications (*63, 73*). In brief, with treatment with DMSO or 2 μM PPZ for 4 h, 5-day-old seedlings (approximately 1 g per biological replicate) cultivated on ½- strength MS medium under dark conditions were fixed using 1% formaldehyde under vacuum for 15 minutes, followed by grinding into fine powder in liquid nitrogen. Chromatin was sonicated to obtain fragments of approximately 300 base pairs using a Bioruptor sonicator, utilizing a 30/30-second on/off cycle (27 total on cycles) at the high setting. Immunoprecipitation was performed overnight at 4°C using 1 μl of anti GFP antibody (Abcam, Cat. No. ab290). ChIP-qPCR analysis was conducted with three biological replicates, and the results were quantified as a percentage of input DNA, following the guidelines provided in the Champion ChIP-qPCR user manual (SABioscience). Primers used for constructing are listed in table S4.

For ChIP-seq, approximately 2 g of seedlings was utilized, and the ChIPed DNA was purified using the MinElute PCR purification kit (Qiagen, Cat. No. 28004). Libraries were constructed using 1-2 ng of ChIPed DNA with the VAHTS Universal DNA Library Prep Kit for Illumina V3 (Vazyme Biotech, Cat. No. ND607), VAHTS DNA Adapters set3–set6 for Illumina (Vazyme Biotech, Cat. No. N805), and VAHTS DNA Clean Beads (Vazyme Biotech, Cat. No. N411-02), following the manufacturer’s protocol. High-throughput sequencing was performed on the Illumina NovaSeq platform (sequencing method: NovaSeq-PE150). The ChIP-seq of BRIP1/2 and BRD1/2/13 using 14-day-old seedlings under long-day conditions has been described (*47, 48*).

ChIP-seq data analysis was conducted following established protocols (*63*). Briefly, the raw data underwent trimming using fastp software, with the parameters set as follows: “-g -q 5 -u 50 -n 15 -l 150”. The resulting clean data was then aligned to the A. thaliana reference genome (TAIR10) using Bowtie 2 with default settings(*74*). Only reads that mapped perfectly and uniquely were retained for subsequent analysis. Peak calling was performed using MACS 2.074 with the following parameters: “gsize = 119,667,750, bw = 300, q = 0.05, nomodel, extsize = 200.” The aligned reads were converted to wiggle (wig) format, and bigwig files were generated using bamCoverage with the options “-bs 10” and “-normalizeUsing RPKM (reads per kilobase per million)” from the deepTools (*75*) software suite. The resulting data were imported into the Integrative Genomics Viewer (IGV) for visualization. Only peaks that were identified in both biological replicates, meeting the threshold of irreproducible discovery rate ≥ 0.05, were considered for further analysis. To annotate the peaks to genes, ChIPseeker (*76*) was employed with default settings, with the requirement of 2 kb upstream and downstream of the transcription start site (TSS). ComputeMatrix and plotProfile (*75*) tools were utilized to compare the mean occupancy density of BRM and BZR1 at specific loci, with detailed information provided in the respective figure legends.

To assess read density and evaluate the correlation between different ChIP-seq samples, we conducted Person correlation analysis. The read density was examined across the combined set of binding sites from all ChIP experiments using the multiBigwig-Summary function available in deepTools (*75*), employing a bin size of 1,000. The Person correlation heatmap was generated using the PlotCorrelation function within deepTools (*75*). To investigate peak overlaps, we employed the Bedtools intersect function, which enabled the identification of common regions between different sets of peaks.

### ATAC-seq assay and data analyses

To isolate protoplasts from 5-day-old Arabidopsis seedlings grown under dark conditions with indicated treatment, approximately 0.5 g of plant tissue was collected and cut into small pieces. Subsequently, the chopped tissue was treated with 5 ml of Enzyme solution, composed of 20 mM MES (pH 5.7), 1.5% cellulase R10, 0.4% macerozyme R10, 0.4 M mannitol, 10 mM CaCl_2_, 3 mM β-mercaptoethanol, and 0.1% BSA, following a previously described method (*77*). Protoplasts were then counted using a hemacytometer under a microscope, and approximately 40,000 protoplasts were used for the isolation of nuclei. The nuclei were obtained by treating the protoplasts with 5 ml of lysis buffer, consisting of 1× PBS (pH 7.5), 0.5% Triton X-100, and 1× Complete Protease Inhibitor Cocktail (Roche). After isolation, the crude nuclei were subjected to three washes with Nuclei Extraction Buffer (1× PBS (pH 7.5), 0.25 M Sucrose, 1 mM PMSF, 1 mM β-mercaptoethanol, 0.5% Triton X-100, 1× Complete Protease Inhibitor Cocktail (Roche)), followed by a single wash with Tris-Mg buffer (10 mM Tris-HCl (pH 8.0), 5 mM MgCl_2_). The purified nuclei were then incubated with Tn5 transposome and tagmentation buffer at 37 °C for 30 min (Vazyme Biotech, Cat. No. TD501). Following tagmentation, DNA was purified using the MinElute PCR purification kit (Qiagen, Cat. No. 28004) and subsequently amplified for 9 cycles using the TruePrepTM DNA Library Prep Kit V2 for Illumina® (Vazyme Biotech, Cat. No. TD501). The number of PCR cycles was determined according to previously published methods (*59*). Index primers from the TruePrepTM Index Kit V2 for Illumina® (Vazyme Biotech, Cat. No. TD202) were used for library amplification. Amplified libraries were then purified using VAHTS DNA Clean Beads (Vazyme Biotech, Cat. No. N411-02). Two biological replicates were performed for each sample. High-throughput sequencing was carried out on the Illumina NovaSeq platform using the NovaSeq-PE150 sequencing method. The ATAC-seq of Col and *brm-1* in Supplementary Fig. 8 using 14-day-old seedlings under short-day conditions.

ATAC-seq data analyses were conducted following previously published methods with some modifications (*59*). Briefly, the raw data underwent trimming using fastp, with the adapter sequence set as “CTGTCTCTTATACACATCT”. The resulting clean data was then mapped to the A. thaliana reference genome (TAIR10) using Bowtie 2 with default settings. To eliminate unmapped and organelle reads, the online tool “Filter BAM datasets” was employed with the parameter “mapping quality ≥30, !Mt, !Pt”. Duplicated reads were removed using the online tool “MarkDuplicates” with default settings. Peak calling, peak annotation, bamCoverage, and visualization in IGV followed the methods employed for ChIP-seq data analyses. Differential DNA accessibility between mutant and wild-type samples was determined using DiffBind (*78*) with default settings. ComputeMatrix and plotProfile (*75*) were utilized to compare the mean DNA accessibility density of mutants and wild-type at defined genomic loci, with detailed information provided in the corresponding figure legends.

### Gene ontology analysis

GO analysis for enriched biological processes was performed with the online tools (https://metascape.org/) with default settings, and plotted at online tools (http://www.bioinformatics.com.cn/)

### Accession numbers

Accession numbers of genes reported in this study include: *AT1G75080* (BZR1), *AT2G46020* (BRM), *AT1G21700* (SWI3C), *AT3G01890* (SWP73A), *AT3G03460* (BRIP1), *AT5G17510* (BRIP2), *AT1G20670* (BRD1), *AT1G76380* (BRD2), *AT5G55040* (BRD13), *AT3G23250* (MYB15), *AT5G52050* (DTX50), *AT5G39860* (PRE1), *AT3G15540* (IAA19), *AT4G34760* (SAUR50)

## Acknowledgements

We thank the Arabidopsis Biological Resource Center (ABRC) for seeds of T-DNA insertion lines, Hongwei Xue (Shanghai Jiao Tong University) for providing *bri1-5* transgenic seeds, Prof. Junxian He (The Chinese University of Hong Kong) for providing *bzr1-1D* seeds, Prof. Jia Li (Guangzhou University) for providing *bri1-701* seeds.

## Author contributions

C.L. conceived the project. T.Z. performed most of the experiments. C.W. generated the *bzr1-1D brm-1* double mutant. T.Z., Y.Y., and J.Z. conducted bioinformatics analysis. T.Z., C.W., Y.Y., J.Z., Y.C., Z.W., and C.L. analyzed data. C.L. wrote the manuscript.

## Data and materials availability

The ChIP-seq, ATAC-seq, and RNA-seq datasets have been deposited in the Gene Expression Omnibus under accession no. GSE233416 and GSE233415, respectively. The BRD1, BRD2, and BRD13 ChIP-seq data were downloaded from GEO under accession no. GSE161595. BRIP1 and BRIP2 ChIP-seq data were downloaded from GEO under accession no. GSE142369. The H3K27me3 ChIP-seq data were downloaded from GEO under accession no. GSE145387. The H3K4me3 ChIP-seq data were downloaded from GEO under accession no. GSE183987. The Pol II and H3K4me2 ChIP-seq data were downloaded from DDBJ databases under the accession number DRR235325 and DRA010413. The H3K36me3 ChIP-seq data were downloaded from GEO under accession no. GSE205112. The H3K9ac, H3K27ac, H4K5ac, H4K8ac, H4K12ac and H4K16ac ChIP-seq data were downloaded from GEO under accession no. GSE183987.

---

**Fig. S1.**
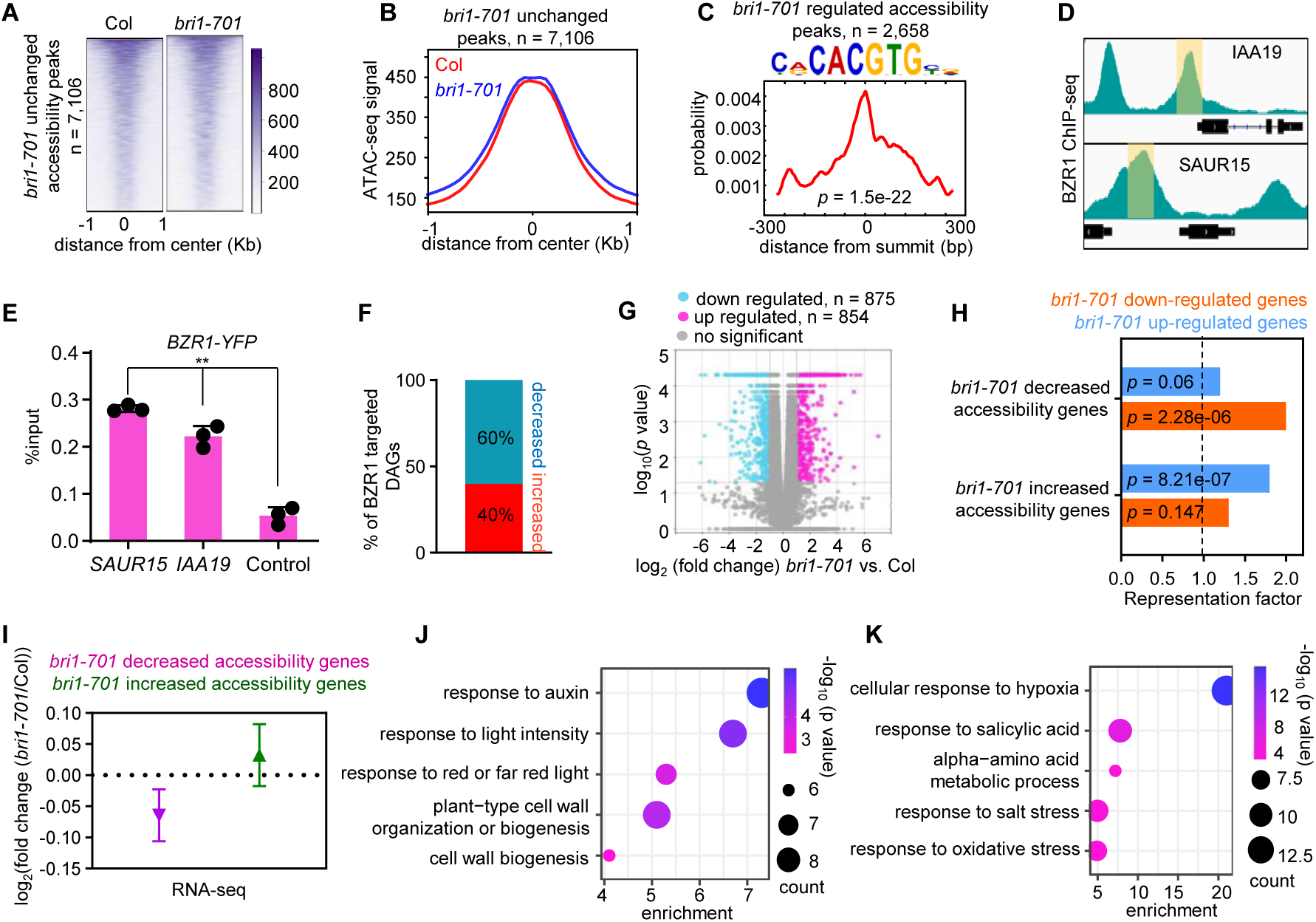
Genome-wide changes of chromatin accessibility and RNA transcription in the loss of BR signaling. (**A**), (**B**) Heatmap (A) and metagene plots (B) reflecting the ATAC-seq signals over the unchanged chromatin accessibility sites for the indicated ATAC-seq experiments. (**C**) The G-Box element is significantly enriched in *bri1-701* regulated accessibility peaks. (**D**) IGV view of BZR1 ChIP-seq at the known BZR1-targeted genes. The black diagrams underneath indicate gene structure. The y-axis scales represent shifted merged MACS2 tag counts for every 10-bp window. (**E**) Validation of the occupancy at the selected sites by ChIP-qPCR in the indicated transgenic plants. Mean ± s.d. from three biological replicates. Statistical significance was determined by two-tailed Student’s t-test; ** *p* < 0.01. (**F**) The percentage of BZR1 targeted genes showing decreased and increased chromatin accessibility in the *bri1-701* muatnts. (**G**) Volcano plots showing differentially expressed genes (|log_2_(fold change)| ≥ 1) in the *bri1-701* mutants, determined by RNA-seq. (**H**) Overlap analysis of genes showing down-regulated and up-regulated in chromatin accessibility and RNA expression in the *bri1-701* mutants. The x axis represents the observed/expected score. The *p* values were calculated by hypergeometric tests. (**I**) Box plot depicts the log_2_ (fold change) in RNA-seq for *bri1-701* decreased chromatin accessibility genes and *bri1-701* increased chromatin accessibility genes. (**J**) Gene ontology analysis of genes showing down-regulated in chromatin accessibility and genes expression in the *bri1-701* mutants. (**K**) Gene ontology analysis of genes showing up-regulated in chromatin accessibility and gene expression in the *bri1-701* mutants.

**Fig. S2.**
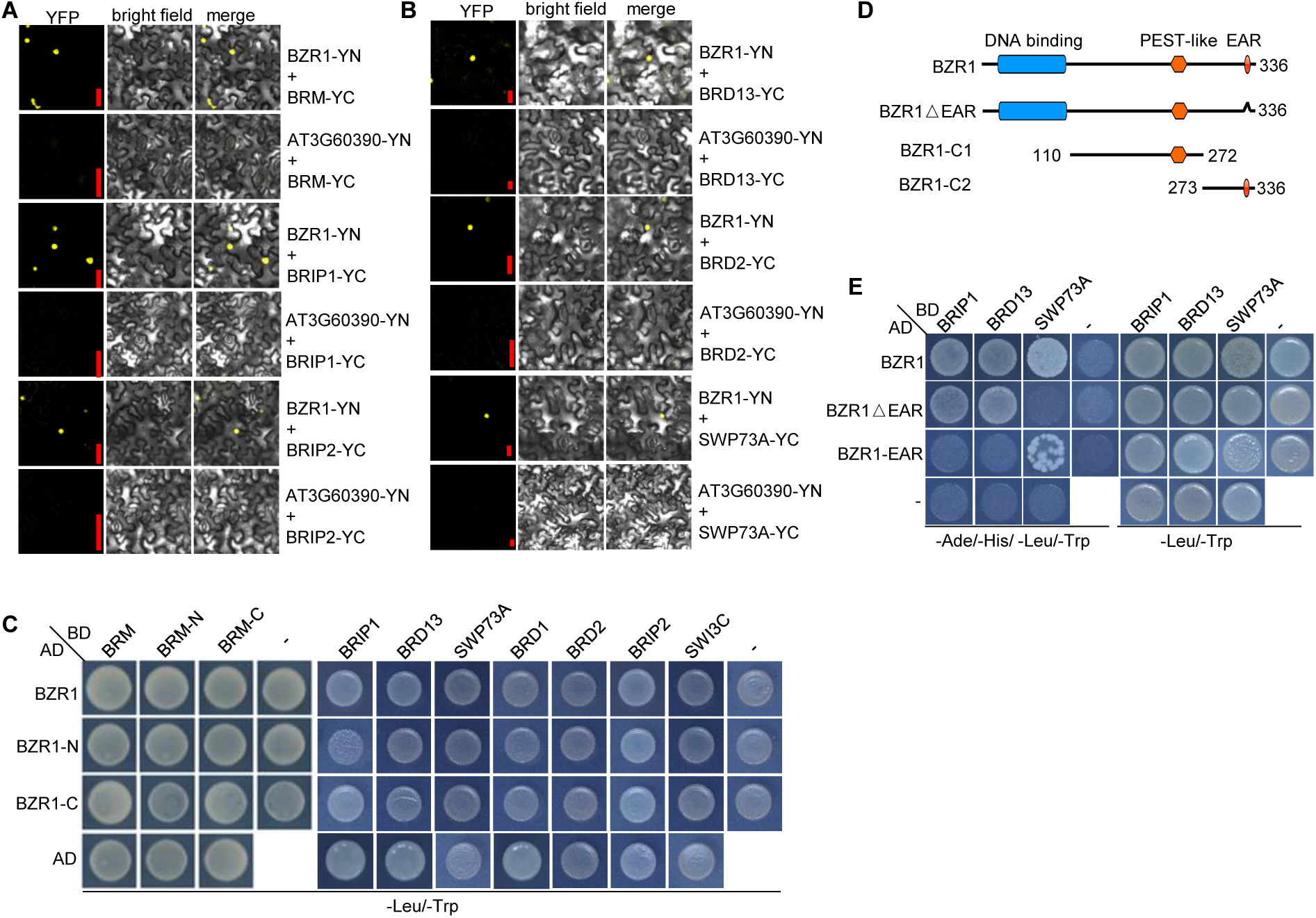
Physical association of BZR1 and BAS complex. (**A**), (**B**) BiFC showing that BZR1 interact with BRM and core members of BAS complex. An unrelated nuclear protein encoded by *AT3G60390* was used as a negative control. error bar = 20 μm. (**C**) Yeast two-hybrid assays to examine BZR1 interact with BRM and core members of BAS complex. Yeast cells transformed with the indicated plasmids were plated onto quadruple dropout (Selective) (SC-Leu, -Trp) medium. AD, Activation Domain; BD, Binding Domain. (**D**) Schematic illustration of the BZR1 and its truncated versions. (**E**) Yeast two-hybrid assays to examine EAR domain of BZR interacts with SWP73A. Yeast cells transformed with the indicated plasmids were plated onto quadruple dropout (Selective) (SC-Ade, - His, -Leu, -Trp) medium. AD, Activation Domain; BD, Binding Domain.

**Fig. S3.**
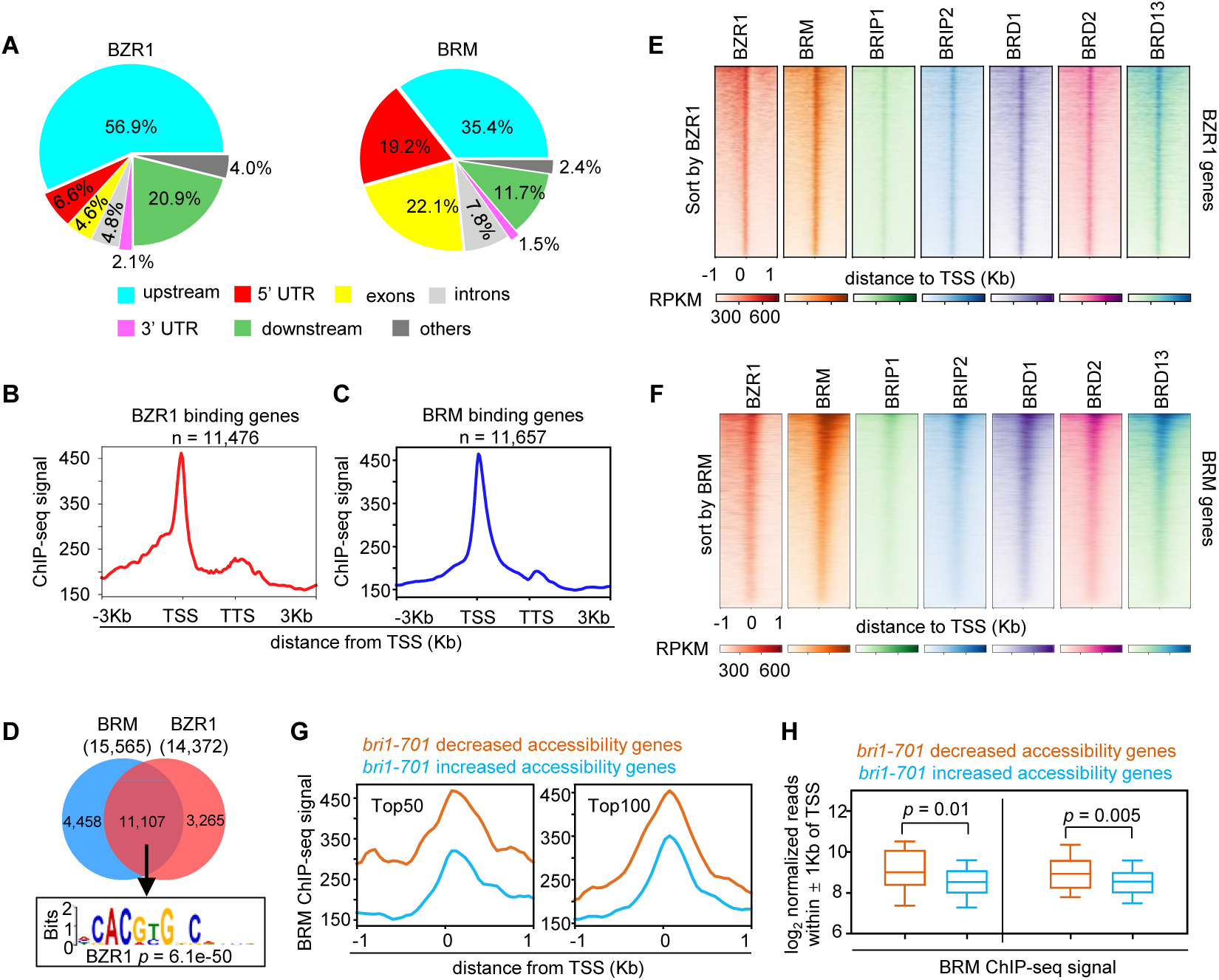
BZR1 and BRM co-occupancy. (**A**) Pie charts showing the distribution of BZR1 and BRM peaks at genic and intergenic regions in the genome. (**B**), (**C**) The average enrichment of BZR1 or BRM over its target genes. Plotting regions were scaled to the same length as follows: 5′ ends (−3.0 kb to transcription starting site (TSS)) and 3′ ends (transcription stop site (TTS) to downstream 3.0 kb), and the gene body was scaled to 2.0 kb. (**D**) The G-Box has a significant enrichment in the BRM and BZR1 overlapped MACS-called peaks. (**E**) Heatmap representations of ChIP–seq of BZR1, BRM, BRIP1/2, and BRD1/2/3. Rank order is from highest to lowest BZR1 signal. log_2_ enrichment was normalized to reads per genome coverage. Read counts per gene were averaged in 50-nucleotide (nt) bins. (**F**) Heatmap representations of ChIP– seq of BZR1, BRM, BRIP1/2, and BRD1/2/3. Rank order is from highest to lowest BRM signal. log_2_ enrichment was normalized to reads per genome coverage. Read counts per gene were averaged in 50-nucleotide (nt) bins. (**G**) Metagene plots displaying the ChIP-seq signals of BRM at the TSS of 50 genes (top 50) or 100 genes (top 100) showing decreased or increased accessibility in the *bri1-701* mutants. (**H**) Box plots displaying read counts for the BRM ChIP-seq dataat *bri1-701* decreased or increased accessibility genes. Reads were summed ± 1 kb from the TSS. Significance analysis was determined by two tailed Mann-Whitney U test.

**Fig. S4.**
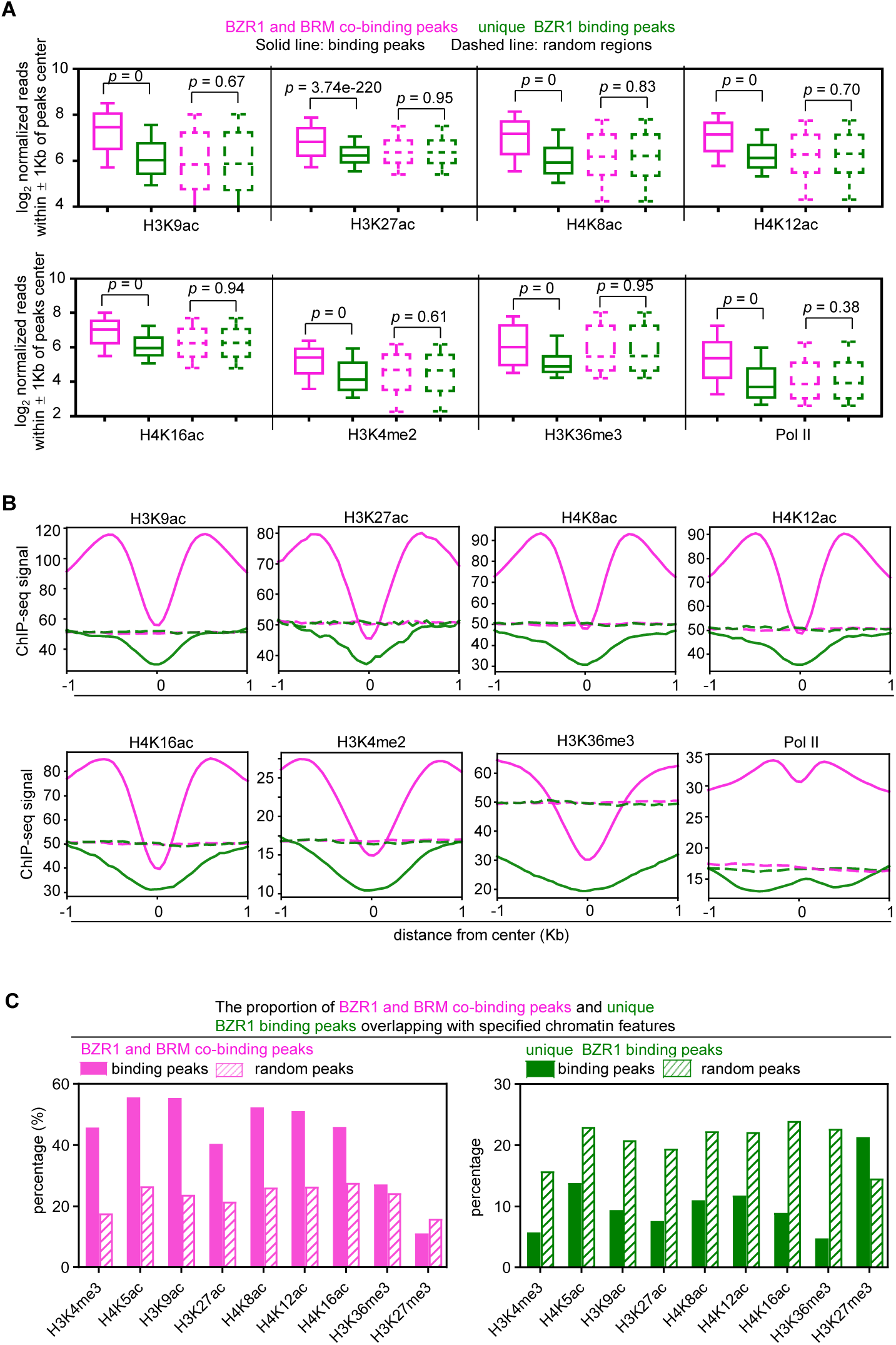
Metagene plots displaying the ChIP-seq signals of different histone modifications at BZR1 and BRM co-binding peaks or unique BZR1 binding peaks. (**A**) Box plots displaying read counts at BZR1 and BRM co-binding peaks or unique BZR1 binding peaks for the H3K9ac, H3K27ac, H4K8ac, H4K12ac, H4K16ac, H3K4me2, H3K36me3, and Pol II ChIP-seq data. Reads were summed ± 1 kb from the peak center. Significance analysis was determined by two tailed Mann-Whitney U test. (**B**) Metagene plots displaying the ChIP-seq signals of H3K9ac, H3K27ac, H4K8ac, H4K12ac, H4K16ac, H3K4me2, H3K36me3, and Pol II at BZR1 and BRM co-binding peaks or unique BZR1 binding peaks. (**C**) The proportion of at BZR1 and BRM co binding peaks or unique BZR1 binding genes overlapping with specified chromatin features.

**Fig. S5.**
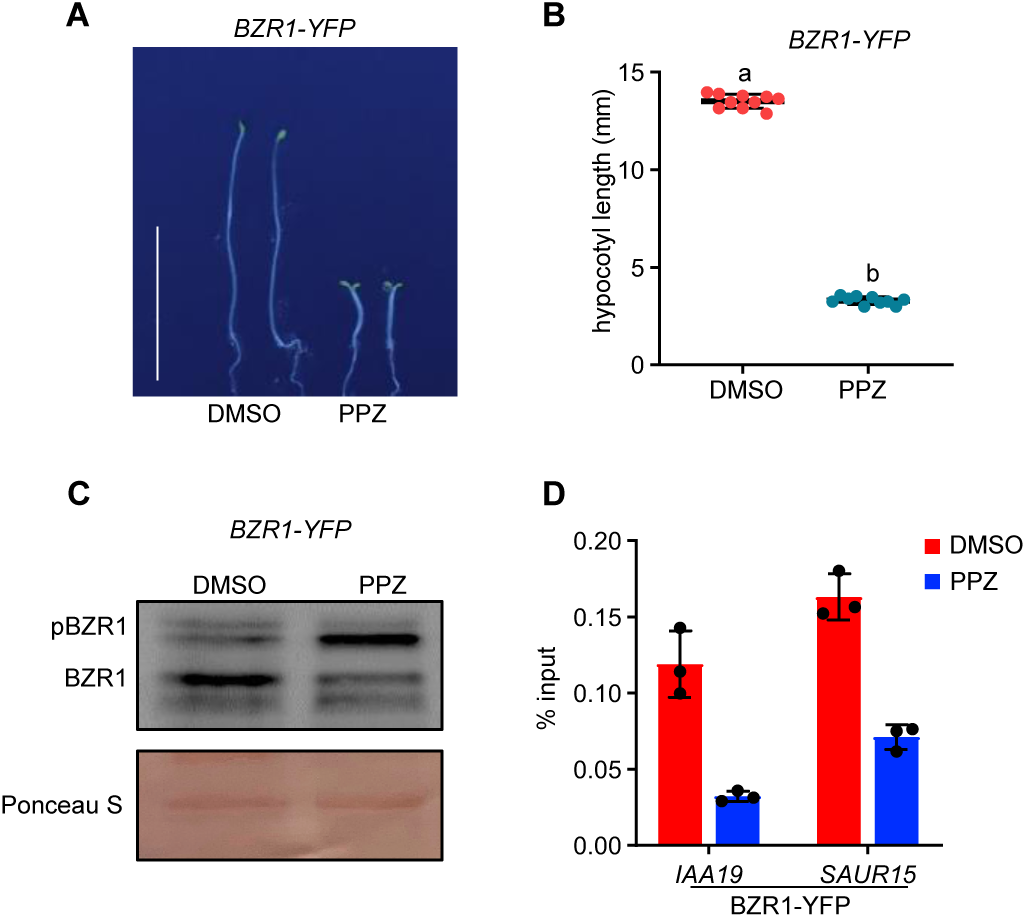
The occupancy of BZR1 is decreased with PPZ treatment. (**A**), (**B**) Hypocotyl elongation phenotypes of *BZR1-YFP* seedlings were shown in dark for 5 days on 1/2 MS medium with DMSO or 2 µM PPZ. The hypocotyl lengths of the indicated genotypes were measured and are shown in **B**. Data are means ± SD. n=10. Scale bars, 10 mm. (**C**) Immunoblot analysis showing the relative protein levels of BZR1 with treatment of DMSO or 2µM PPZ. (**D**) Validation of BZR1 enrichment at *IAA19* and *SAUR15* loci by ChIP–qPCR with treatment of DMSO or 2µM PPZ.

**Fig. S6.**
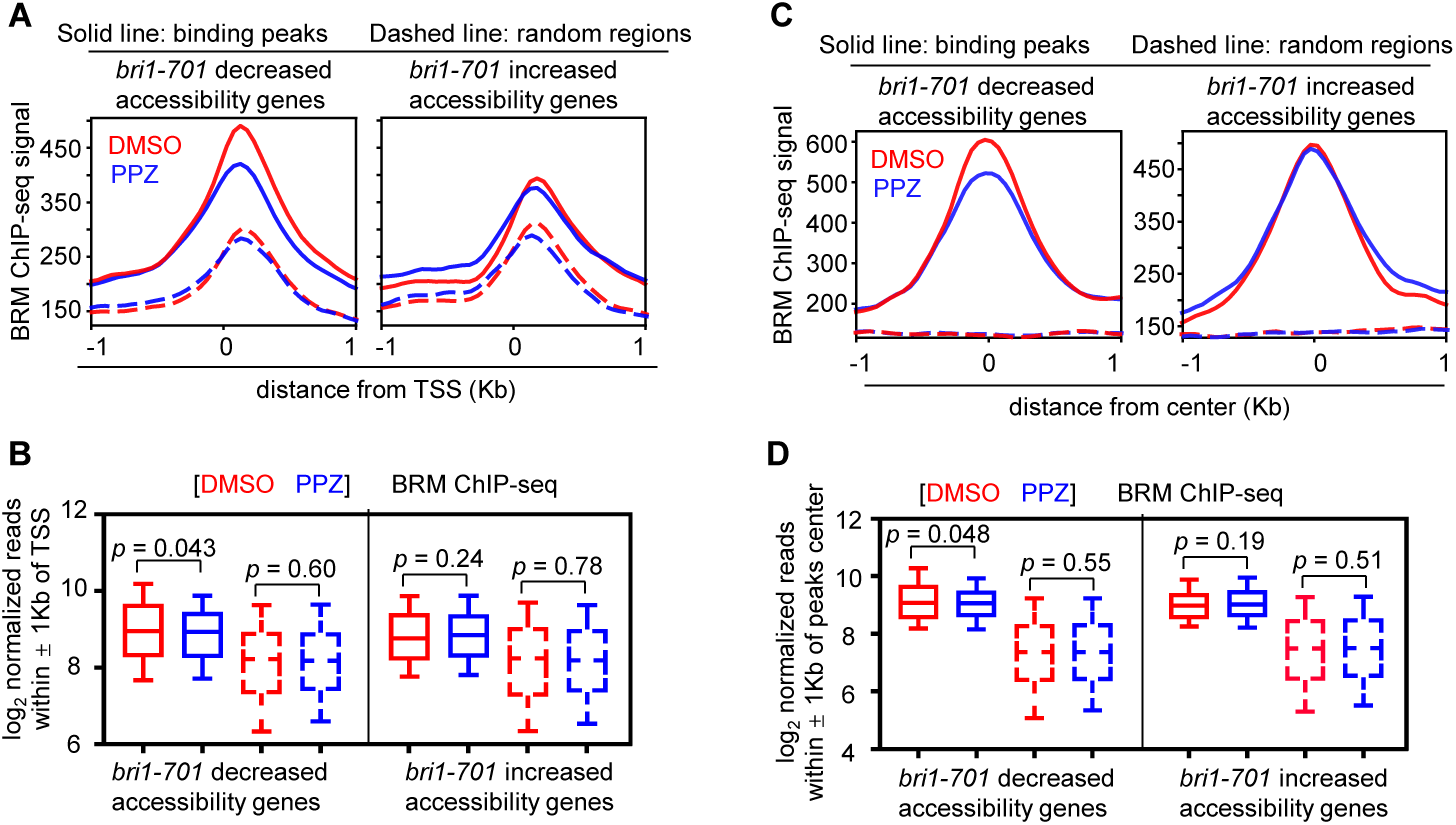
BR enhances BRM enrichment signal at *bri1-701* decreased accessibility sites. (**A**) Metagene plots displaying the ChIP-seq signals of BRM at the TSS of *bri1-701* decreased or increased accessibility genes. (**B**) Box plots displaying read counts for the BRM ChIP-seq data at *bri1-701* decreased or increased accessibility genes. Reads were summed ± 1 kb from the TSS. Significance analysis was determined by two tailed Mann-Whitney U test. (**C**) Metagene plots displaying the ChIP-seq signals of BRM binding peaks at *bri1-701* decreased or increased accessibility genes. (**D**) Box plots displaying read counts for the BRM ChIP-seq data at *bri1-701* decreased or increased accessibility genes. Reads were summed ± 1 kb from the peaks center. Significance analysis was determined by two tailed Mann-Whitney U test.

**Fig. S7.**
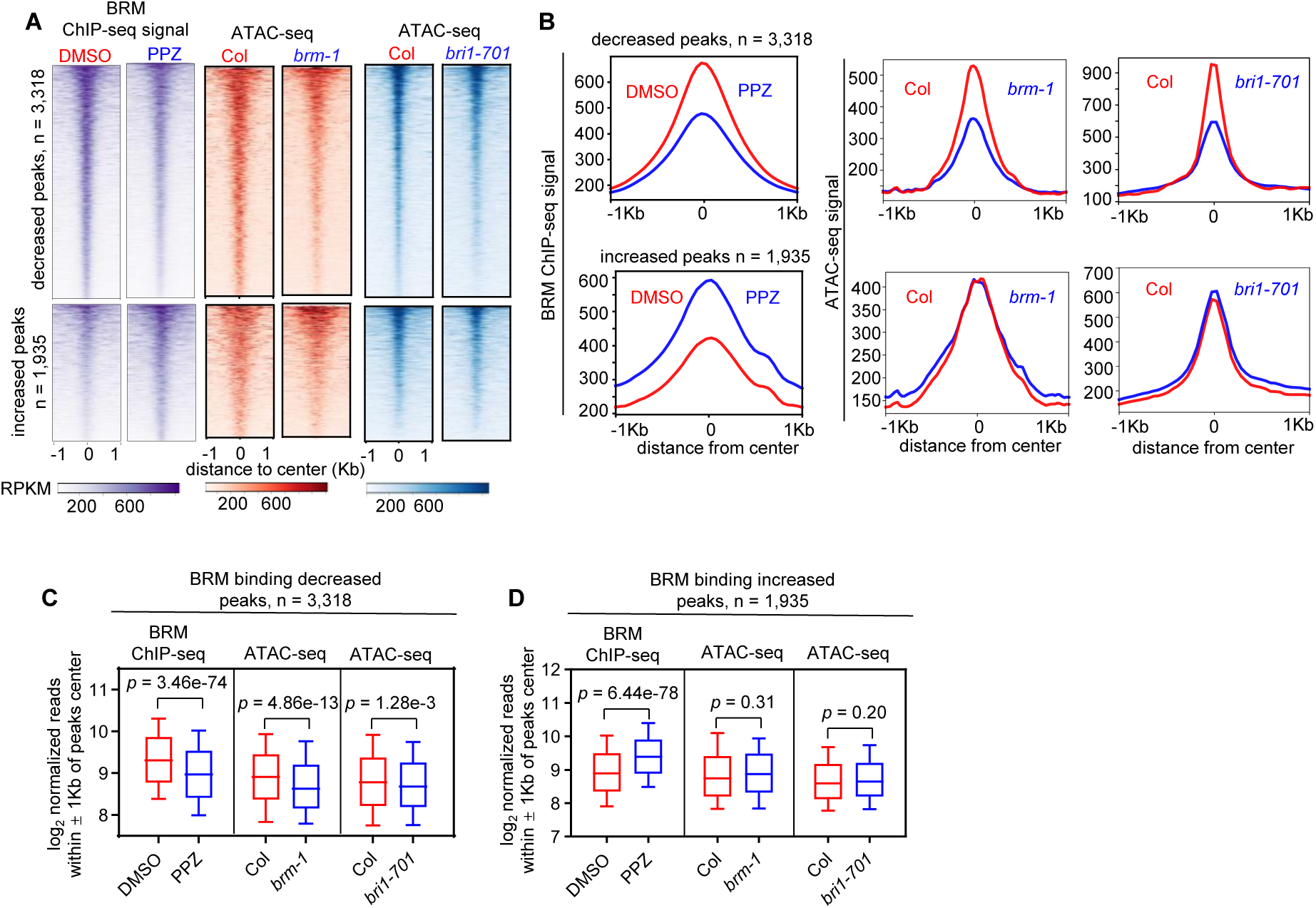
*brm-1* and *bri1-701* showed a similar decline in accessibility at BRM binding decreased sites. (**A**), (**B**) Heatmap (A) and metagene plots (B) reflecting the ChIP-seq signals and ATAC-seq signal at the decreased, or increased BRM binding sites. (**C**), (**D**) Box plots displaying read counts for the BRM ChIP-seq or ATAC-seq data at BRM binding decreased or BRM binding increased peaks. Reads were summed ± 1 Kb from the peaks center. Significance analysis was determined by two tailed Mann-Whitney U test.

**Fig. S8.**
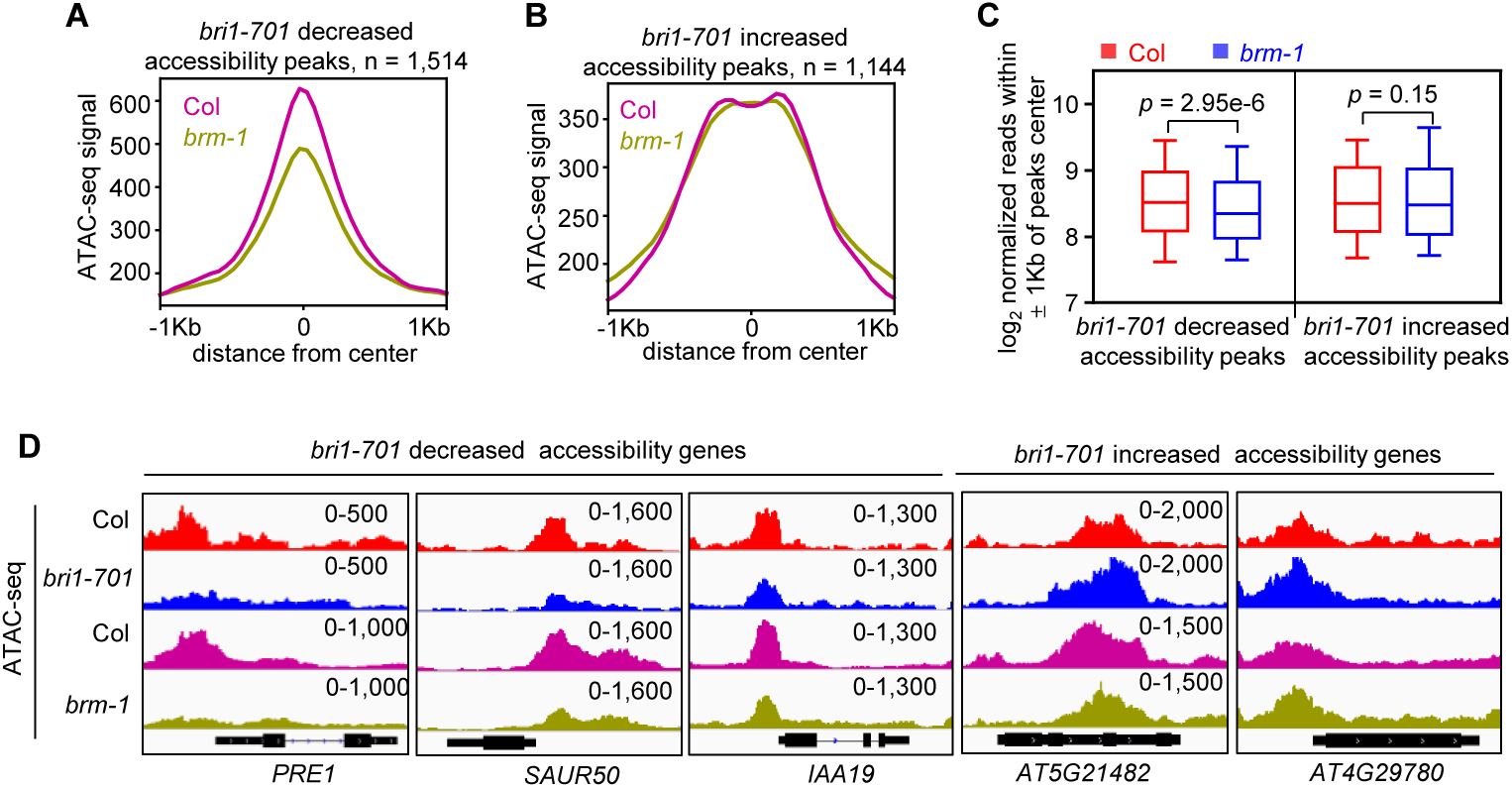
The loss of BRM results in a decline in chromatin accessibility at *bri1-701* decreased accessibility sites. (**A**, **B**) Metagene plots reflecting the ATAC-seq signals over the *bri1-701* decreased or increased chromatin accessibility sites for the indicated ATAC-seq experiments. (**C**) Box plot displaying read counts over the decreased or increased chromatin accessibility sites for the indicated ATAC-seq experiments. Reads were summed ± 1 kb from the peaks center. Significance analysis was determined by two tailed Mann-Whitney U test. (**D**) Examples of ATAC-seq tracks at representative loci in the Col, *bri1-701* and *brm-1* mutants.

**Fig. S9.**
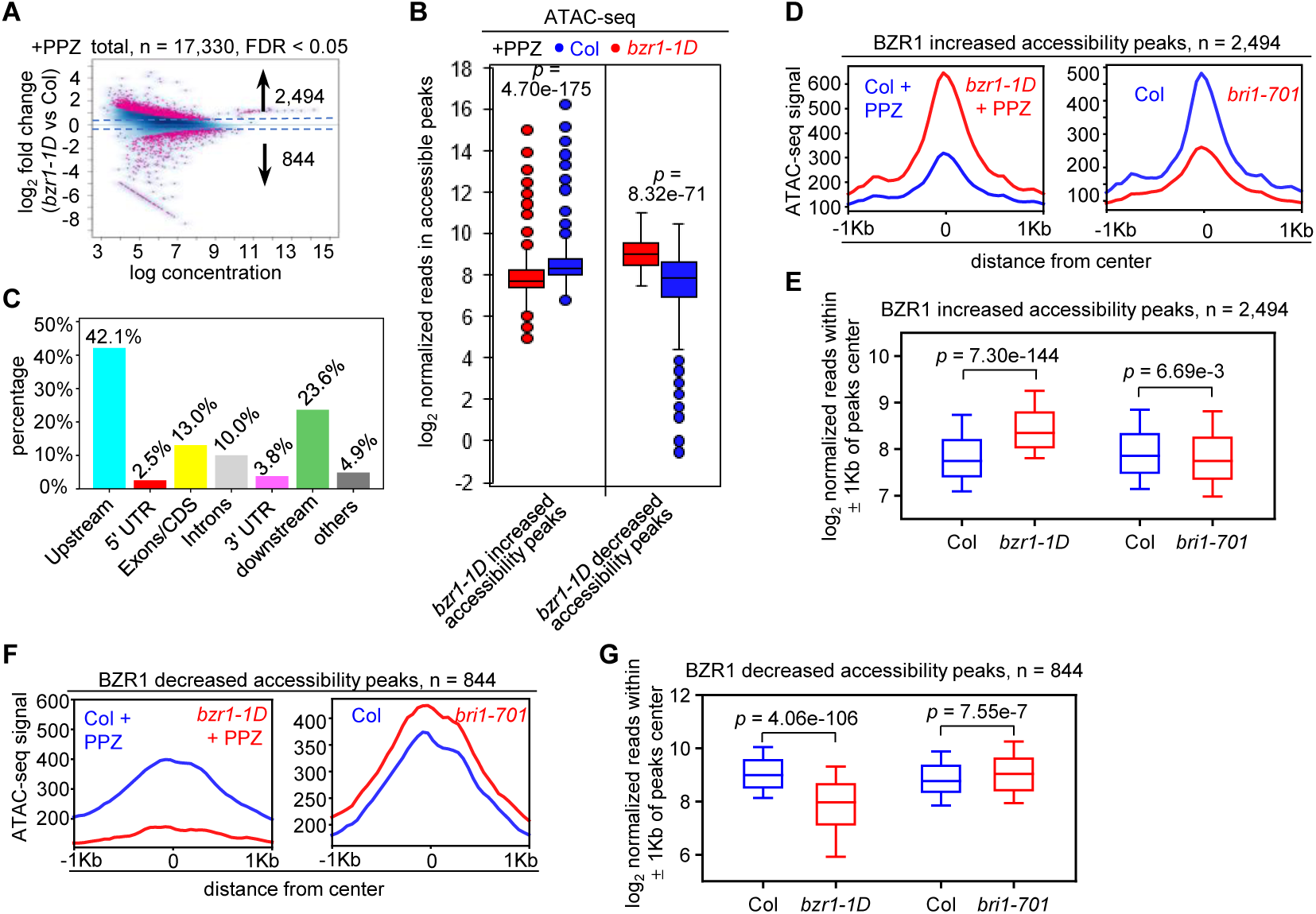
*bzr1-1D* rescues the changes of chromatin accessibility with treatment of PPZ. (**A**) Scatter plot showing fold-change (|log2 fold change| ≥ 0.4) of accessible peaks between WT and *bzr1-1D*. Blue dots, stable peaks; pink dots, differential peaks. The numbers of differentially accessible peaks (increased or decreased) according to FDR are indicated. (**B**) Box plots showing read counts at regions that had increased and decreased accessibility in *bzr1-1D* for the indicated ATAC-seq experiments. Significance analysis was determined by two tailed Mann-Whitney U test. (**C**) Bar chart showing the distribution of changed chromatin accessible peaks in the *bzr1-1D* mutants at genic and intergenic regions in the genome. (**D**), (**E**) Metagene plots and box plot reflecting the ATAC-seq signals over the BZR1 increased chromatin accessibility regions for the indicated assays. Significance analysis was determined by two tailed Mann-Whitney U test. (**F**), (**G**) Metagene plots and box plot reflecting the ATAC-seq signals over the BZR1 decreased chromatin accessibility regions for the indicated assays. Significance analysis was determined by two tailed Mann Whitney U test.

**Fig. S10.**
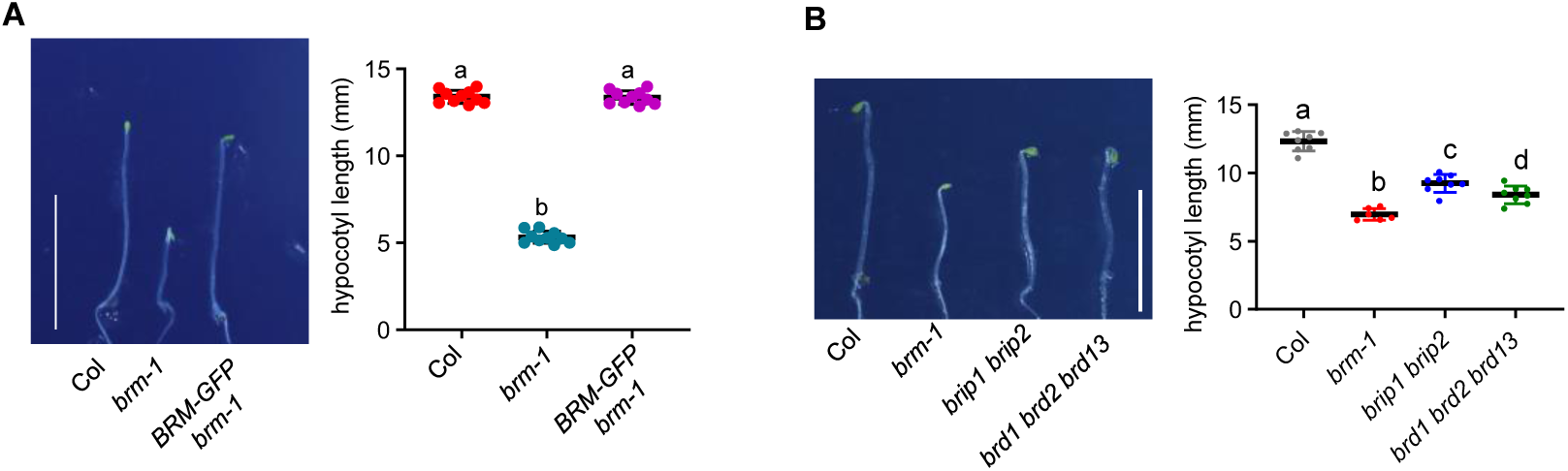
Loss of BAS complex reduces hypocotyl elongation. (**A**) The hypocotyl elongation phenotype of indicated lines grown in dark for five days. The error bars in the right graph indicate the s.d. (n = 10 plants). Lowercase letters indicate statistical significance determined by the Student’s t test. Scale bars, 10 mm. (**B**) The loss of core subunits of MAS inhibits the promotion of hypocotyl elongation. Seedlings were grown in the dark for five days. Lowercase letters indicate statistical significance determined by the Student’s t test. Scale bar, 10 mm.

